# Phosphatidylserine Receptors Enhance SARS-CoV-2 Infection: AXL as a Therapeutic Target for COVID-19

**DOI:** 10.1101/2021.06.15.448419

**Authors:** Dana Bohan, Hanora Van Ert, Natalie Ruggio, Kai J. Rogers, Mohammad Badreddine, José A. Aguilar Briseño, Roberth Anthony Rojas Chavez, Boning Gao, Tomasz Stokowy, Eleni Christakou, David Micklem, Gro Gausdal, Hillel Haim, John Minna, James B. Lorens, Wendy Maury

## Abstract

Phosphatidylserine (PS) receptors are PS binding proteins that mediate uptake of apoptotic bodies. Many enveloped viruses utilize this PS/PS receptor mechanism to adhere to and internalize into the endosomal compartment of cells and this is termed apoptotic mimicry. For viruses that have a mechanism(s) of endosomal escape, apoptotic mimicry is a productive route of virus entry. We evaluated if PS receptors serve as cell surface receptors for SARS-CoV-2 and found that the PS receptors, AXL, TIM-1 and TIM-4, facilitated virus infection when low concentrations of the SARS-CoV-2 cognate receptor, ACE2, was present. Consistent with the established mechanism of PS receptor utilization by other viruses, PS liposomes competed with SARS-CoV-2 for binding and entry. We demonstrated that this PS receptor enhances SARS-CoV-2 binding to and infection of an array of human lung cell lines and is an under-appreciated but potentially important host factor facilitating SARS-CoV-2 entry.

## INTRODUCTION

Severe Acute Respiratory Syndrome Coronavirus 2 (SARS-CoV-2) emerged in late 2019 and quickly spread around the world, resulting in the current public health pandemic. SARS-CoV-2 is a beta coronavirus of the sarbecovirus subgenus and is closely related to SARS-CoV, the agent responsible for an epidemic in 2003. SARS-CoV-2 is effectively transmitted between humans and has infected more than 178 million individuals and caused more than 3.88 million deaths worldwide as of June 23, 2021 (WHO). Fortunately, a herculean scientific effort has resulted in the development of SARS-CoV-2 vaccines which have been shown to be efficacious, potentially stemming the pandemic. Nonetheless, in combination with vaccines, continued development of efficacious antivirals is needed, as outbreaks continue in under-vaccinated regions and severe disease is not eradicated following vaccination. Towards this goal, a more comprehensive understanding of SARS-CoV-2 interactions with host cells will be required.

SARS-CoV-2 entry into cells is mediated by the viral spike glycoprotein (S) binding to Angiotensin Converting Enzyme 2 (ACE2) (1-3). The S1 subunit of S binds to ACE2 while S2 mediates membrane fusion. Cleavage at the S1/S2 junction occurs during virus egress from producer cells by the host protease furin which facilitates S1 binding to ACE2. A second site termed S2’ is also cleaved by the host proteases. Cleavage by TMPRSS2 at the cell surface promotes fusion of the viral and host plasma membranes (2, 4). Alternatively, SARS-CoV-2 virions can be internalized via clathrin-mediated endocytosis after ACE2 binding, wherein host cathepsins (especially cathepsin L) proteolytically cleave S2’ (3, 5-7). Adherence factors that enhance virion binding and increase infectivity have also been identified, namely neuropilin 1 and heparan sulfate (8, 9).

Binding and internalization of a variety of different enveloped viruses occurs through virion associated phosphatidylserine (PS) binding to PS receptors. Members of the TIM family (TIM-1 and TIM-4) bind PS directly while another family of PS receptors, the TAM tyrosine kinase receptor family (TYRO3, AXL and MERTK), bind PS indirectly through the adaptor proteins Gas6 and Protein S. These PS receptor mediate binding and internalization of a wide range of viruses, including filoviruses, alphaviruses, and flaviviruses (10-13). TIM-1, TIM-4, and AXL appear to be the most efficacious at mediating viral entry given their prevalent use among enveloped viruses (14-16). Once virions are within the endosome, events that result in virion fusion with cellular membranes are virus specific, with filoviruses requiring viral glycoprotein processing followed by interactions with Niemann Pick C1 protein (NPC1) to initiate fusion, whereas flaviviruses rely on endosomal acidification driving glycoprotein conformational changes which mediate fusion (17, 18).

Given that PS receptors mediate entry of other enveloped viruses through interactions with viral membrane PS, we assessed the role of PS receptors on SARS-CoV-2 infection and the mechanism of interaction. We found that plasma membrane-expressed PS receptors by themselves do not result in productive SARS-CoV-2 infection; however, these receptors enhance infection when low levels of ACE2 are expressed. Our findings indicate that these receptors synergize with ACE2 to mediate SARS-CoV-2 entry through PS-dependent interactions. These data are in contrast to the conclusions drawn by another report stating that AXL interacts directly with the SARS-CoV-2 spike protein (19). Appreciation of this route of entry provides an additional pathway that could be therapeutically targeted to inhibit virus entry and subsequent infection.

## RESULTS

### PS receptors enhance ACE2-dependent SARS-CoV-2 infection

The ability of TIM and TAM family PS receptors to support SARS-CoV-2 infection were initially examined in transfected HEK 293T cells. Wild-type HEK 293T cells do not express significant amounts of ACE2 or PS receptors and are poorly permissive to SARS-CoV-2 infection (3, 20). Expression plasmids encoding ACE2 and/or the PS receptors, AXL or TIM-1, were transfected, resulting in expression of these receptors on the surface of the transfected cells (**S1A-B**). Dual transfection did not alter expression of ACE2 or PS receptors relative to single transfection. The PS receptors AXL and TIM-1 by themselves did not facilitate infection of SARS-CoV-2 or VSV pseudovirions bearing SARS-CoV-2 spike (VSV/Spike) (**Fig. 1A, S1C**). However, when low levels of ACE2 (50 to 250 ng of plasmid) were co-expressed with either AXL or TIM-1, the combinations enhanced infection over that observed with ACE2 alone. TIM-1-enhanced ACE2-dependent recombinant VSV/Spike (rVSV/Spike) (21) infection, and enhancement occurred over a wider range of ACE2 concentrations than AXL-enhanced ACE2-dependent infection, with AXL consistently enhancing infection when 250 ng of ACE2 plasmid was transfected (**Fig. 1B-C, S1C**). The more limited ability of AXL to synergize was not due to limiting Gas6 in the media as the addition of Gas6 to media did not enhance the synergy. At higher concentrations of ACE2 plasmid, PS receptor enhancement of infectivity was reduced and, with transfection of 1 μg of ACE2 plasmid, no PS receptor enhancement was observed. Thus, only when ACE2 is limiting on the cell surface do PS receptors facilitate infection. Consistent with a role for PS receptors in SARS-CoV-2 entry, we observed enhanced virion attachment to cells when PS receptors were expressed (**Fig. 1D**). In addition to viral load assessments, supernatants from SARS-CoV-2 infected HEK 293T cultures were evaluated for production of infectious virions at 48 hpi by TCID_50_ assays in Vero E6 cells that express TMPRSS2. Low levels (50 ng) of ACE2 transfection increased virion production and this was enhanced by co-expression of TIM-1 (**Fig. 1E**). Consistent with the viral load findings, AXL co-expression was not effective at enhancing production of infectious virus when this low level of AXL was transfected. Other PS receptors, TIM-4, TYRO3, and MerTK were examined for their ability of enhance infection. TIM-4 enhanced ACE2-dependent entry of VSV/Spike in a manner similar to TIM-1; however, TYRO3 and MerTK of the TAM family did not mediate increased entry, despite detectable levels of plasma membrane expression after transfection (**Fig. 1F; S1A-B; S1D**). The synergy between PS receptors and ACE2 was specific for SARS-CoV-2 as infection with VSV-luciferase pseudovirions bearing Lassa virus GP was not affected by expression of these receptors (**S1E**). These studies indicate that PS receptors synergize with low levels of ACE2 to enhance SARS-CoV-2 infection. Further, these data provide evidence that VSV/Spike pseudovirions serve as a useful BSL2 surrogate for SARS-CoV-2 entry events as others have shown (22-24).

**Figure 1:**
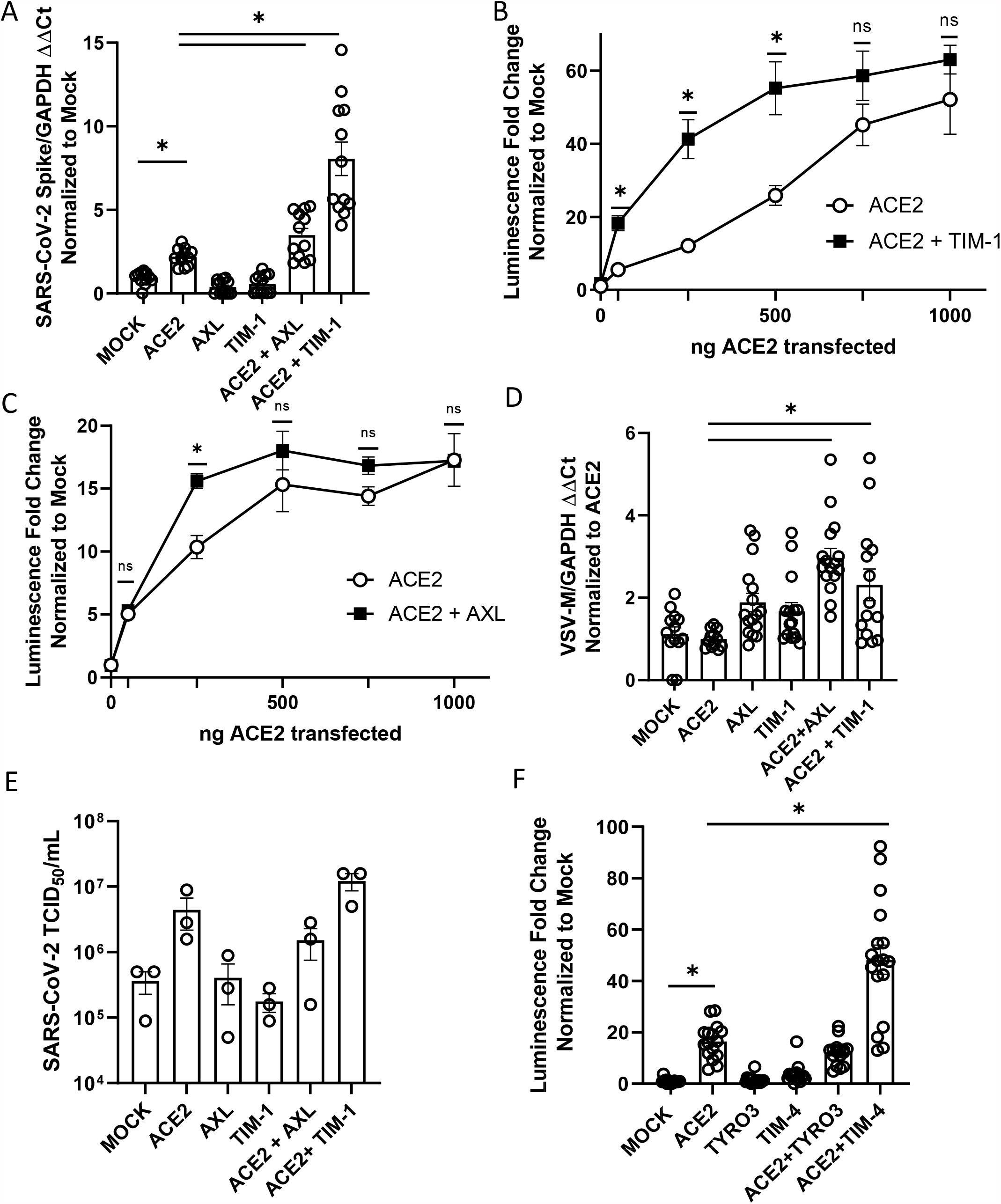
PS receptors synergize with ACE2, enhancing SARS-CoV-2 infection of HEK 293T cells. **A) C**ells transfected with expression PS receptor plasmids, AXL or TIM-1, with or without 50 ng of ACE2 and infected 48 hours later with SARS-CoV-2 (MOI = 0.5). Viral loads were determined 24 hours following infection. **B-C)** PS receptors, TIM-1 **(B)** and AXL **(C)**, enhance rVSV-luciferase/Spike infection at low concentrations of ACE2 are transfected. **D)** Virus binding of cells transfected with PS receptor plasmids with or without 50 ng of ACE2. rVSV/Spike was bound to transfected cells at 48 hpi and bound virus was measured via RT-qPCR. **E)** Supernatants from SARS-CoV-2 infected (MOI = 0.5) transfected HEK 293T cells were titered 48 hpi on Vero E6-TMPRSS2 and TCID_50_ calculated by Spearman-Karber equation. These studies were performed with transfection of 50 ng of ACE2 plasmid. **F)** HEK 293T cells transfected with expression PS receptor plasmids, TYRO3 or TIM-4, with or without 50 ng of ACE2 and infected 48 hours later with SARS-CoV-2 (MOI = 0.5). Viral loads were determined 24 hours following infection. Data shown are pooled from at least 3 independent experiments (**A, B, C, D, E, F**). Data represented as means ± SEM. Student’s t-test **(A**,**E)** and multiple t-test **(B**,**C)**, One-Way ANOVA with multiple comparisons **(D&F)**; asterisks represent p < 0.05.

### PS receptors bind to virion associated PS, not the SARS-CoV-2 Spike protein

We took several different approaches to examine the mechanism by which PS receptors interact with SARS-CoV-2. In the context of other viral pathogens, PS receptors are known to bind to virion membrane associated PS and mediate endosomal internalization of virions, shuttling virus to cognate endosomal receptors. Liposomes composed of PS compete for virion binding to PS receptors (25). To test this with SARS-CoV-2, increasing concentrations of PS or phosphatidylcholine (PC) liposomes were evaluated for their ability to compete with virus for PS binding sites in ACE2 + AXL or ACE2 + TIM-1 transfected HEK 293T cells. PS liposomes effectively blocked VSV/Spike entry, whereas PC liposomes were significantly less effective (**Fig. 2A-B**). We also assessed the activity of a TIM-1 mutant, ND115DN which has a disrupted TIM-1 PS binding pocket, for its ability to facilitate VSV/Spike entry. This TIM-1 mutant did not synergize with ACE2, indicating that the TIM-1 PS binding pocket is critical for this enhanced activity (**Fig. 2C**) (12, 20).

**Figure 2:**
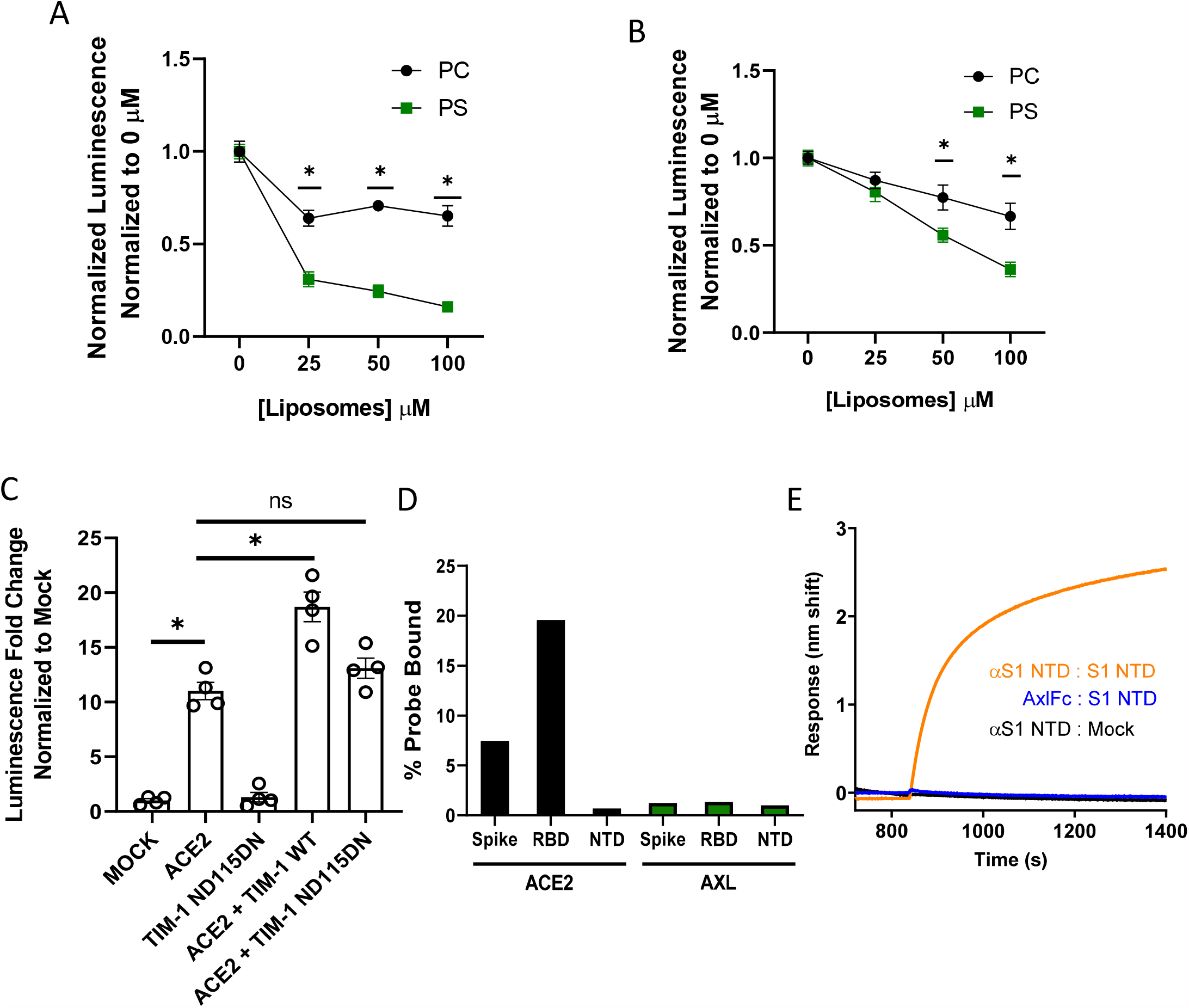
PS receptors interact with SARS-CoV-2 by binding to PS. **A-B)** PS liposomes interfere with rVSV-luciferase/Spike infection. HEK 293T cells transfected with TIM-1 plasmid and 50 ng of ACE2 plasmid (**A**) or AXL plasmid and 50 ng of ACE2 plasmid (**B**) were infected with rVSV-luciferase/Spike in the presence of increasing concentrations of PS or PC liposomes and assessed for luciferase activity at 24 hours following infection. **C)** HEK 293T cells were transfected with WT or PS binding pocket mutant TIM-1 plasmids with or without 50 ng of ACE2 expressing plasmid and infected 48 hours later with rVSV-luciferase/Spike pseudovirions. Luminescence fold change were compared to mock transfected lysates that were set to a value of 1. **D)** AXL is unable to directly interact with purified, soluble SARS-CoV-2 spike/Fc. HEK 293T cells transfected with AXL or ACE2 were incubated with soluble spike protein (S1/S2)-Fc, S1 RBD-Fc or S1 NTD-Fc and subsequently incubated with an Alexa 647 secondary. Spike protein binding was detected by flow cytometry. **E)** AXL does not bind to the NTD of SARS-CoV-2 spike. Biolayer interferometry association curves show that immobilized AXL-Fc fails to interact with purified NTD of spike. Data are pooled from at least 3 independent experiments (**A, B**) or are representative of at least 3 experiments (**C, D, E**). Data represented as means ± SEM. Multiple t-test (**A, B**), One-way ANOVA with multiple comparisons (**C**); asterisks represent p < 0.05.

Others have reported that the N-terminal domain of SARS-CoV-2 spike directly binds to AXL and is important for AXL-mediated entry of SARS-CoV-2 (19). To assess spike/AXL interactions, purified soluble full-length SARS-CoV-2 spike-Fc, spike receptor binding domain-Fc (RBD) or spike N-terminal domain-Fc (NTD) was incubated with HEK 293T cells transiently expressing AXL. Flow cytometry was used to detect spike proteins bound to AXL. Parallel ACE2 binding to soluble SARS-CoV-2 spike served as a positive control. As the spike NTD-Fc was not expected to bind to ACE2, an ELISA confirmed the ability of a conformationally dependent α-spike NTD monoclonal antibody to bind NTD-Fc, suggesting that NTD-Fc was in its native conformation (**S2B**). The full-length spike-Fc and the RBD-Fc bound to ACE2, but no interactions were detected between any of the purified spike proteins and AXL (**Fig. 2D**) despite evidence of robust AXL surface expression on transfected HEK 293T cells (**S2A**) and the equivalent levels of detection of the purified proteins via ELISAs (**S2C**). Biolayer interferometry studies confirmed and extended our findings that recombinant AXL does not bind to purified NTD, whereas interaction with the α-spike NTD monoclonal antibody was readily detected (**Fig. 2E**). Thus, using two complementary approaches, we were unable to demonstrate direct interactions of AXL with spike. In total, our studies are consistent with PS receptors interacting with SARS-CoV-2 virions through the well-established mechanism of virion-associated PS binding to TIM-1 and AXL.

### Redundant routes of virus entry: endosomal vs. plasma membrane mediated infection

ACE2-dependent coronaviruses enter cells through two different routes: 1) An endosomal route of virus uptake that requires low pH-dependent cathepsin L processing of spike, and 2) A plasma membrane route that is dependent upon TMPRSS2 cleavage of spike (3, 26). Others have reported that TMPRSS2-dependent entry is preferentially utilized by the virus when this serine protease is expressed (27). We examined the route of virus entry at play when ACE2, PS receptors and/or TMPRSS2 was expressed.

Initially, we assessed enhancement conferred by increasing amounts of TMPRSS2 plasmid on ACE2-dependent infection. As anticipated, we found that low levels (10 ng) of TMPRSS2 expressing plasmid enhanced VSV/Spike pseudovirion entry in HEK 293T cells co-transfected with 50 ng of ACE2 expressing plasmid (**Fig. 3A**). However, at higher concentrations of TMPRSS2 plasmid, TMPRSS2 did not enhance infection, perhaps due to aberrant protease activity. Inverting these variables, when 10 ng of TMPRSS2 plasmid was transfected in the presence of low concentrations of ACE2, TMPRSS2 enhanced virus infection (**Fig. 3B**). At high concentrations of ACE2, virus entry became TMPRSS2-independent in a manner similar to the effects of the PS receptors. Taken with Fig. 1B and C, these studies indicate that the PS receptors and TMPRSS2 can facilitate ACE2-dependent virus infection when ACE2 is limiting, but with increasing ACE2 concentrations the infections become independent of these entry factors. This may be related to effects of soluble ACE2 on entry (28).

**Figure 3:**
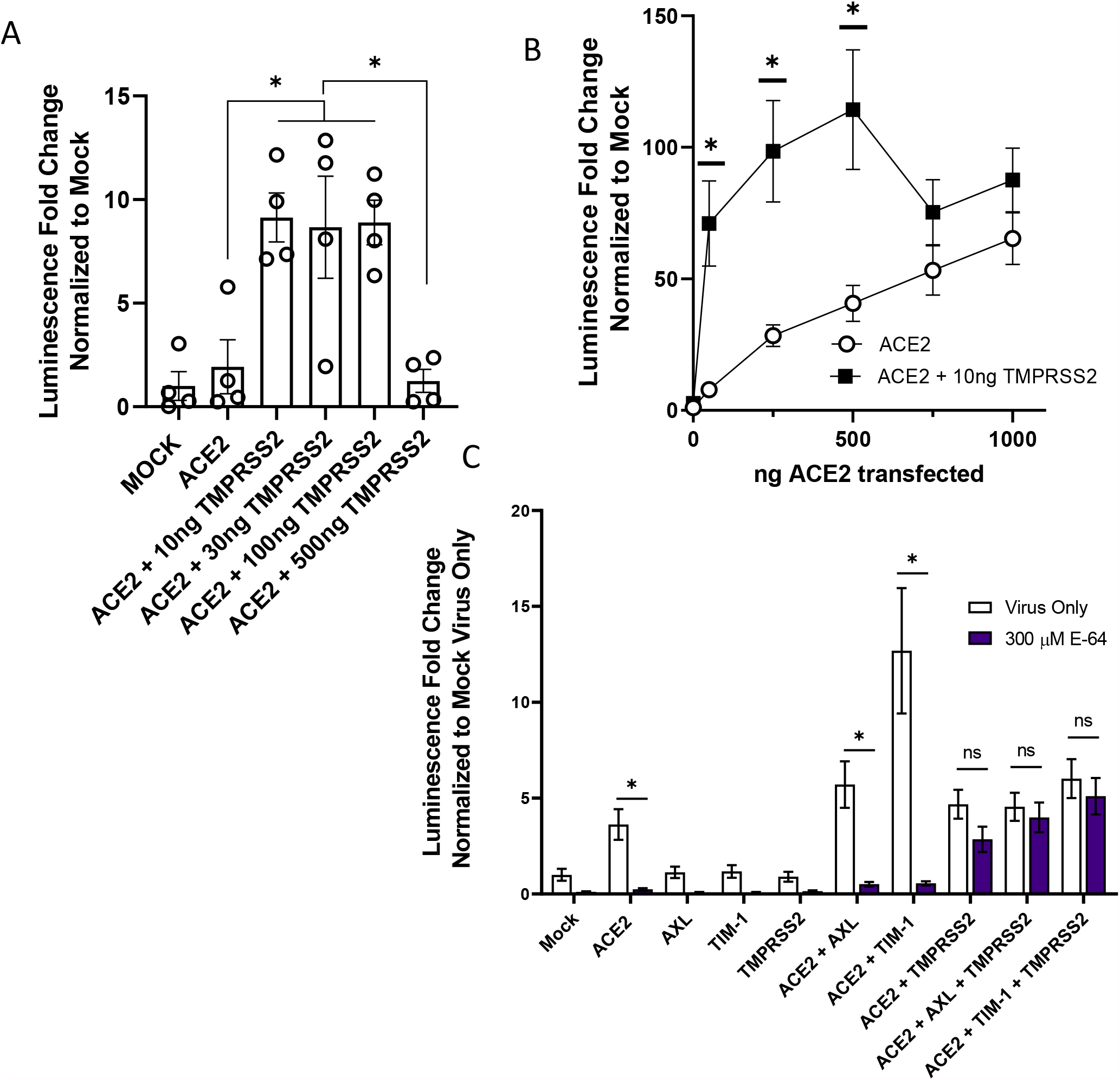
The route of SARS-CoV-2 entry is altered by TMPRSS2 expression. **A)** HEK 293T cells were transfected with ACE2 and TMPRSS2 as noted and infected at 48 h with VSV-luciferase/Spike. At 24 hpi, luminescence activity was determined. Findings are shown relative to empty vector (Mock) transfected cells. Panel depicts one representative experiment. Students t-tests. **B)** TMPRSS2 expression enhances rVSV-luciferase/Spike entry at low levels of ACE2 expression. HEK 293T cells were transfected as indicated and pseudovirion entry assessed by measuring luminescence activity at 24 hpi. **C)** Transfected HEK 293T cells were transfected and infected with VSV-luciferase/Spike at 48 h in the presence or absence of E-64 (300 μM). Luciferase activity was determined 24 hpi. Data are pooled from at least 3 independent experiments (**B, C**) or are representative of at least 3 experiments (**A**). Data represented as means ± SEM. Student’s T-tests **(A)** Multiple t-tests (**B**), Two-way ANOVA with row-wise multiple comparisons (**C**); asterisks represent p < 0.05.

To evaluate how PS receptors and/or TMPRSS2 expression would alter the route of ACE2-dependent infection, HEK 293T cells were transfected PS receptors as before and infected with VSV/Spike in the presence or absence of the cysteine protease inhibitor, E-64, that blocks endosomal cathepsin activity. Non-toxic levels of E-64 were effective at blocking ACE2-dependent infection (**Fig. 3C, S3**), indicating that virions were entering these cells in a cysteine protease-dependent manner, likely through the endosomal compartment. The enhancement of virus entry conferred by the combination of PS receptors and ACE2 was also inhibited by E-64, providing evidence that this is the route of virion uptake that is enhanced by PS receptors. These findings are consistent with earlier reports that PS receptors mediate cargo internalization into the endosomal compartment (12, 13, 29). In cells that expressed ACE2 and TMPRSS2, virus entry was no longer sensitive to E-64 as previously reported (6, 27). VSV/Spike entry in the presence of TMPRSS2, PS receptors, and ACE2, was also insensitive to E-64, suggesting that the TMPRSS2 expression and activity mediates entry at the plasma membrane independent of PS receptor utilization.

### Inhibition of endogenous AXL utilization blocks SARS-CoV-2 entry

We next evaluated the ability of endogenously expressed PS receptors to enhance SARS-CoV-2 in ACE2 positive cells. Vero E6 cells that express ACE2, AXL, and TIM-1 (**S4A**) were initially assessed (20). Notably, a large fraction of ACE2 protein is located intracellularly, suggesting a rich reserve of ACE2 is inaccessible to extracellular virions (**S4B**). Initial studies using PS liposomes confirmed that PS receptors are important for SARS-CoV-2 infection of these cells. Competition studies in Vero E6 cells demonstrated that increasing doses of PS, but not PC, inhibited VSV/Spike infection, similar to our findings in HEK 293T cells (**Fig. 4A**). PS liposomes also significantly reduced SARS-CoV-2 binding to the surface of Vero E6 cells, implicating PS receptors in attachment and subsequent entry of SARS-CoV-2 (**Fig. 4B**). These findings reinforce the importance of either AXL, TIM-1, or both for SARS-CoV-2 entry.

**Figure 4:**
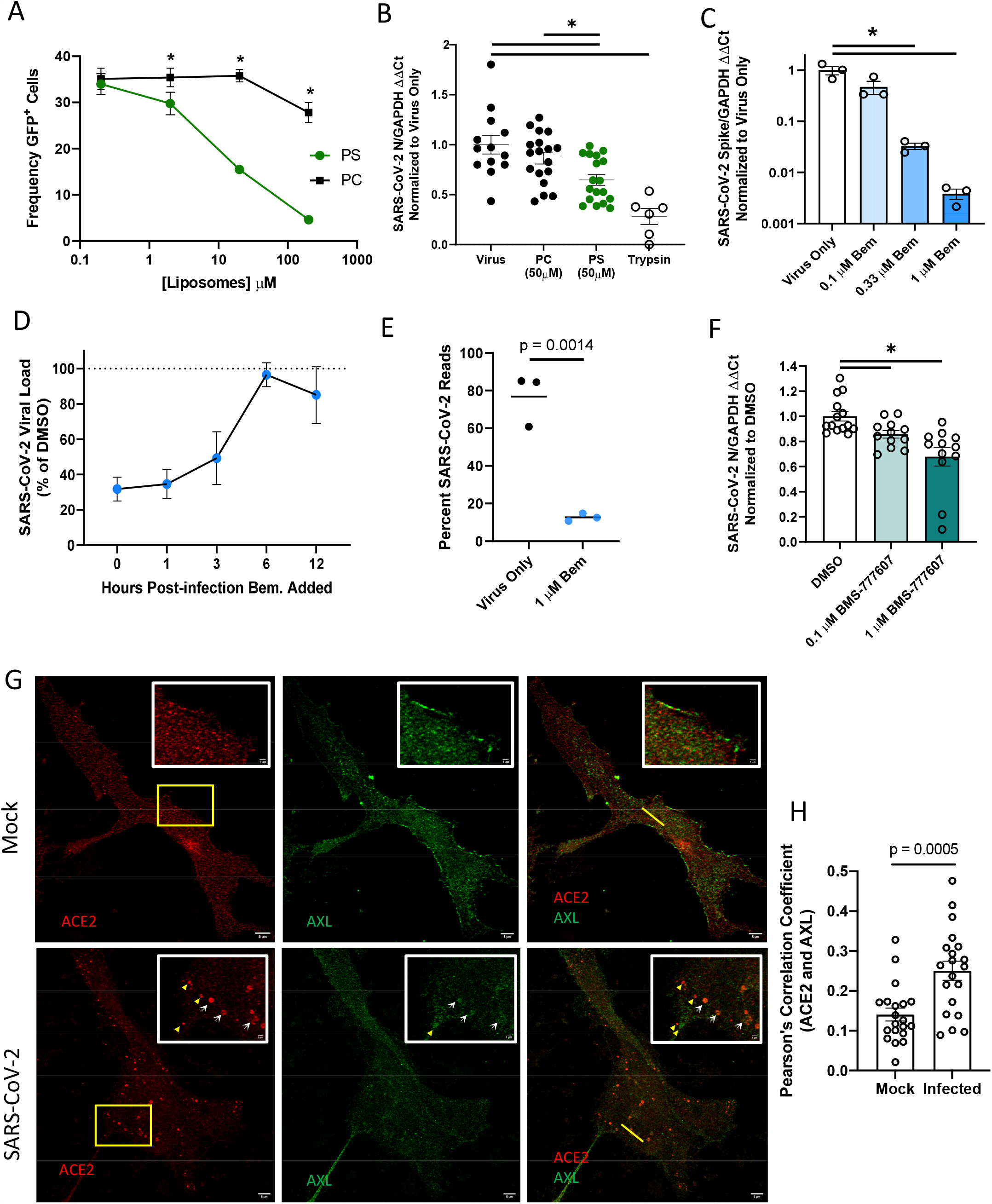
AXL has a prominent role in SARS-CoV-2 entry in Vero E6 cells. **A)** PS liposomes interfere with SARS-CoV-2 pseudovirion entry. Vero E6 cells were treated with PS or PC liposomes and incubated with VSV-GFP/Spike pseudovirions for 24 hours. Entry was detected by GFP fluorescence. **B)** PS liposomes disrupt SARS-CoV-2 binding. Vero E6 cells were incubated with SARS-CoV-2 (MOI = 5) at 10°C for 1 hour, washed extensively, and viral load assessed by RT-qPCR. **C)** AXL signaling inhibitor bemcentinib inhibits SARS-CoV-2 infection in Vero E6 cells. Cells were treated with bemcentinib and infected with SARS-CoV-2 (MOI = 0.01). Viral loads were measured 24 hpi by RT-qPCR. **D)** Bemcentinib inhibition of SARS-CoV-2 infection is most efficacious at early time points during infection. Vero E6 cells were challenged with SARS-CoV-2 (MOI = 0.01) and treated with either the vehicle control or 1 μM bemcentinib at the indicated time. Viral loads were measured 24 hpi by RT-qPCR. **E)** Vero E6 cells were treated with 1 μM bemcentinib, infected with SARS-CoV-2 (MOI = 0.01) and mRNA harvested 18 hpi. mRNA was deep sequenced on an Illumina platform, and viral loads were calculated by alignment to the SARS-CoV-2 genome. **F)** Broad spectrum TAM inhibitor BMS-777607 inhibits SARS-CoV-2 infection in Vero E6 cells. Cells were treated with inhibitor at indicated concentrations for 1 hour, challenged (MOI = 0.01), and viral loads measured 24 hpi by RT-qPCR **G)** STED micrographs showing staining for ACE2 (red) and AXL (green) and merged in Vero E6 cells. Insets are enlarged images from regions highlighted by yellow rectangles. White arrows indicate shared vesicular structures between the two channels. Yellow arrowheads indicate objects that are only seen in one channel. Plot profiles are shown in **S4F**, representing signal intensity along the yellow lines in the merged panels. **H)** Pearson’s correlation coefficients of ACE2 and AXL were calculated for n=20 mock and infected cells (ROI determined by cell borders). Data are pooled from at least 3 independent experiments (**B, D, F**) or are representative of at least 3 experiments (**A, C, G, H**). Data are represented as means ± SEM. Multiple t-tests (**A**) student’s t-test (**B, C, F, H**); asterisks represent p < 0.05.

To assess if AXL was important for infection of Vero E6 cells, the selective AXL kinase inhibitor, bemcentinib, was tested for its ability to block SARS-CoV-2 infection. In a dose dependent manner, bemcentinib profoundly inhibited SARS-CoV-2 virus load and blocked infection of the one-hit VSV/Spike pseudovirion (**Fig. 4C, S4D**). A time-of-addition study indicated that bemcentinib inhibition was most effective at early timepoints during SARS-CoV-2 infection, consistent with a role of AXL in virus entry (**Fig. 4D**). Bemcentinib toxicity was tested on human lung epithelial cells and was nontoxic at the concentrations used (**S4C**). Consistent with an important role for AXL in SARS-CoV-2 infection, RNAseq of infected Vero E6 demonstrated that infection of these cells with a MOI of 0.01 resulted in ∼80% of transcripts composed of viral transcripts 18 hpi (**Fig. 4E**). If these cells were treated with 1 μM bemcentinib at the time of infection, the fraction of viral transcripts dropped precipitously, decreasing to ∼10% of the total reads.

To determine if TIM-1 contributed to SARS-CoV-2 infection of Vero E6 cells, the blocking anti-human TIM-1 monoclonal antibody, ARD5, was evaluated for inhibition of recombinant VSV (rVSV) bearing either Ebola GP (EBOV) or spike. While rVSV/EBOV GP was inhibited by ARD5 as previously reported (12, 30), it had no effect on rVSV/Spike infection (**S4E**). Thus, despite robust expression of both PS receptors, AXL is preferentially utilized for SARS-CoV-2 infection in these cells. While preferential PS receptor utilization has been reported for other pathogens (20), our previous studies indicated that TIM-1 rather than AXL was preferentially used, in contrast to our current observations with SARS-CoV-2. Host factors or virion attributes determining PS receptor preference are currently unexplored. A broad-spectrum TAM inhibitor, BMS777607, modestly reduced virus infection in a dose dependent manner (**Fig. 4F**), discounting the likelihood of off-target effects with bemcentinib.

As the bulk of ACE2 in Vero E6 cells is intracellular (**S4B**), surface expressed-AXL may be facilitating SARS-CoV-2 uptake into the endosomal compartment where proteolytic processing and ACE2 interactions mediate fusion of the viral envelope and cellular membranes. Previous studies with the betacoronavirus responsible for the 2003-2004 outbreak, SARS-CoV, demonstrated that ACE2 is found abundantly in the endosomal compartment, specifically co-localizing with the early endosomal marker EEA1 (31). Further, at 3 hpi, SARS-CoV antigens colocalize with vesicular ACE2 and that ACE2 formed notable vesicular puncta in the infected cells (31). We utilized Stimulated Emission Depletion (STED) microcopy, leveraging the super resolution capabilities of this platform to investigate ACE2 and AXL colocalization in uninfected and infected Vero E6 cells. In uninfected cells, AXL and ACE2 were found on the plasma membrane and intracellularly, but colocalize poorly (**Fig. 4G and H**). However, as shown in the micrographs (white arrows) and the associated fluorescence intensity plot profiles (**S4F** yellow lines highlight selected ROI), ACE2 and AXL demonstrate overlapping localization patterns within cytoplasmic punctate structures. Pearson’s correlation coefficients of infected and uninfected cells calculated for AXL and ACE2 intensity demonstrate a significant increase in colocalization values between this PS receptor and the cognate SARS-CoV-2 receptor in infected cells, relative to mock counterparts. (**Fig. 4H**). These data support the possibility that PS receptors enhance SARS-CoV-2 trafficking into these intracellular puncta where ACE2 is abundant.

### AXL promotes SARS-CoV-2 infection in a range of lung cell lines

In addition to ACE2, many lung cell lines express AXL (19, 32) (**S5A**). We evaluated these lines for their ability to support SARS-CoV-2 infection and whether infection was sensitive to bemcentinib inhibition. The panel of lung cells that were selected included A549 (adenocarcinoma) stably expressing ACE2, HCC1650 (NSCLC), HCC1944 (squamous), H1819 (adenocarcinoma), H2302 (adenocarcinoma) and Calu3 (adenocarcinoma).

These cells were inoculated with SARS-CoV-2 (MOI = 0.5 unless otherwise noted) in the presence or absence of a serial dilution of the AXL inhibitor, bemcentinib, or the cysteine protease inhibitor, E-64. The cell lines A549^ACE2^, H1650, HCC1944, H1819, and HCC2302 readily supported SARS-CoV-2 infection, and viral loads 24 hpi were decreased in a dose-dependent manner, by bemcentinib and E64 (**Fig. 5A-E**). Infectious SARS-CoV-2 present in HCC2302 cell supernatants at 24 and 48 hpi were also markedly decreased by bemcentinib (**Fig. 5F**), demonstrating that bemcentinib treatment reduced production of new infectious virus in a dose dependent manner. Further, at 1 μM of bemcentinib, detectable production of any infectious virus was delayed until 48 hours (L.O.D. = 5 TCID_50_/mL). In H1650 cells, the ability of bemcentinib to inhibit recently emerged SARS-CoV-2 variants of concern (VOC), Alpha (B.1.1.7) and Beta (B.1.351), was evaluated. While the Alpha VOC replicated poorly in these cells, bemcentinib significantly inhibited virus replication of both variants, providing evidence the efficacy of the AXL inhibitor is not influenced by SARS-CoV-2 adaptative changes (**S5B**).

**Figure 5:**
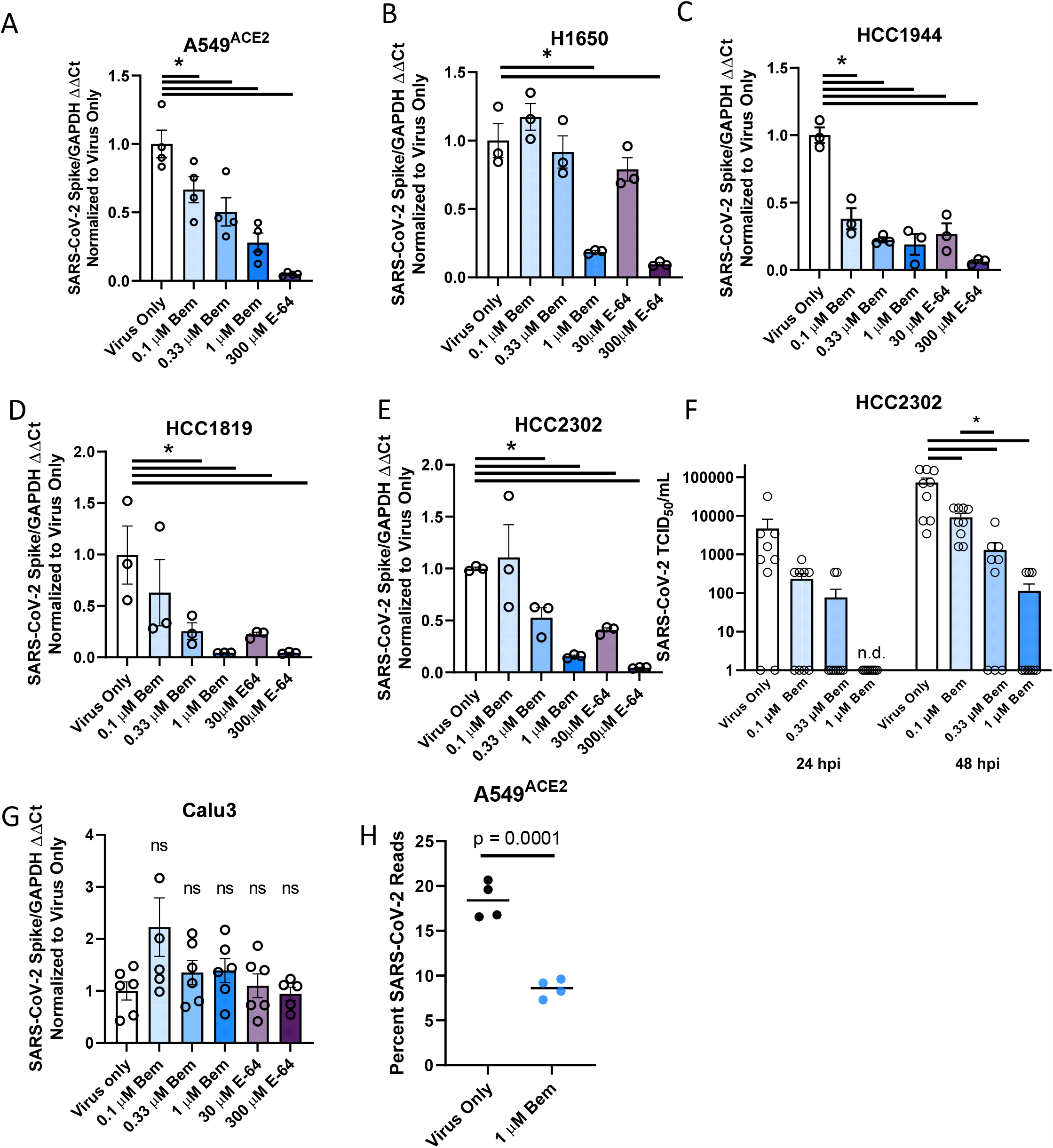
AXL inhibition reduces SARS-CoV-2 infection in human lung cells. **A-F)** SARS-CoV-2 infection is reduced by AXL inhibition in multiple human lung cell lines. In order: A549 ^ACE2^, H1650, HCC1944, H1819, HCC2302, Calu3 were treated with the indicated inhibitors for 1 hour and challenged with SARS-CoV-2 (MOI = 0.5) for 24 hours. Viral load was assessed by RT-qPCR. **G)** HCC2302 cells were treated with bemcentinib at the indicated concentrations for 1 hour and infected with SARS-CoV-2 (MOI = 0.5). Input virus was removed 6 hpi and supernatant was collected at 24 and 48 hpi and titered by TCID_50_ assays on Vero E6-TMPRSS2 cells. TCID_50_ /mL was calculated by the Spearmann-Karber method. **H)** A549 ^ACE2^ were treated with bemcentinib as indicated, infected with SARS-CoV-2 (MOI = 0.5) and mRNA harvested 24 hpi. mRNA was sequenced, and viral loads calculated by alignment to the SARS-CoV-2 genome. Data are pooled from at least 3 independent experiments (**F**) or are representative of at least 3 experiments (**A, B, C, D, E, G**). Data represented as means ± SEM. Student’s t-test; asterisks represent p < 0.05.

SARS-CoV-2 infection of TMPRSS2^+^ Calu-3 cells (**S6A**) was not sensitive to bemcentinib or E-64, again providing evidence that, in this cell line, the route of virus entry was dominated by the TMPRSS2-dependent path, bypassing the use of PS receptors and the endosomal compartment (**Fig. 5G**). These findings stand in contrast to SARS-CoV-2 infection of TMPRSS2^+^ H1650 cells that were markedly bemcentinib and E-64 sensitive and were found to be insensitive to camostat inhibition (**Fig. 5B, S5C**). The paradoxical finding that virus entry into H1650 is sensitive to E64 and bemcentinib despite endogenous TMPRSS2 expression indicates that TMPRSS2-dependent pathways are not always the dominating or default route of SARS-CoV-2 entry and suggests that a more complex balance of events controls which pathway is used. Neither the total amount of cell surface expressed ACE2 nor the intracellular versus extracellular ACE2 ratio appears to determine the route of virus uptake (**S5D**).

RNA sequencing studies confirmed and extended our findings with bemcentinib in A549^ACE2^ cells. At 24 hpi, 20% of the transcripts in A549^ACE2^ cells mapped to the viral genome. Infection in the presence of 1 μM bemcentinib significantly decreased the number of viral transcripts present (**Fig. 5H**). Further analysis of potential qualitative changes in viral transcripts indicated that transcript numbers across the genome were reduced, rather than a reduction of specific subgenomic transcripts.

To directly evaluate the importance of AXL during infection of human lung cells, CRISPR-Cas9 technology was used to knock out (KO) AXL expression in H1650 and HCC2302 cells. H1650 AXL^neg^, a biologically cloned AXL-null line, was evaluated along with bulk AXL KO populations of H1650 and HCC2302, denoted as AXL^low^. The AXL^neg^ clone, which expressed undetectable levels of AXL protein (**Fig. 6A**), supported dramatically lower SARS-CoV-2 virus loads at a range of input MOIs (**Fig. 6B**) and became unresponsive to bemcentinib (**Fig. 6C**), demonstrating an important role for AXL in SARS-CoV-2 infection and indicating that bemcentinib specifically targets AXL. Bulk populations of AXL^low^ H1650 and HCC2302 also supported reduced levels of SARS-CoV-2 infection and were poorly responsive to bemcentinib (**S6A-E**). Taken together, the data presented here demonstrate that SARS-CoV-2 utilizes AXL to enhance virion binding and entry in some lung cell lines, and that this mechanism can be effectively disrupted in human lung cells by small molecule inhibitors and genetic ablation of AXL.

**Figure 6:**
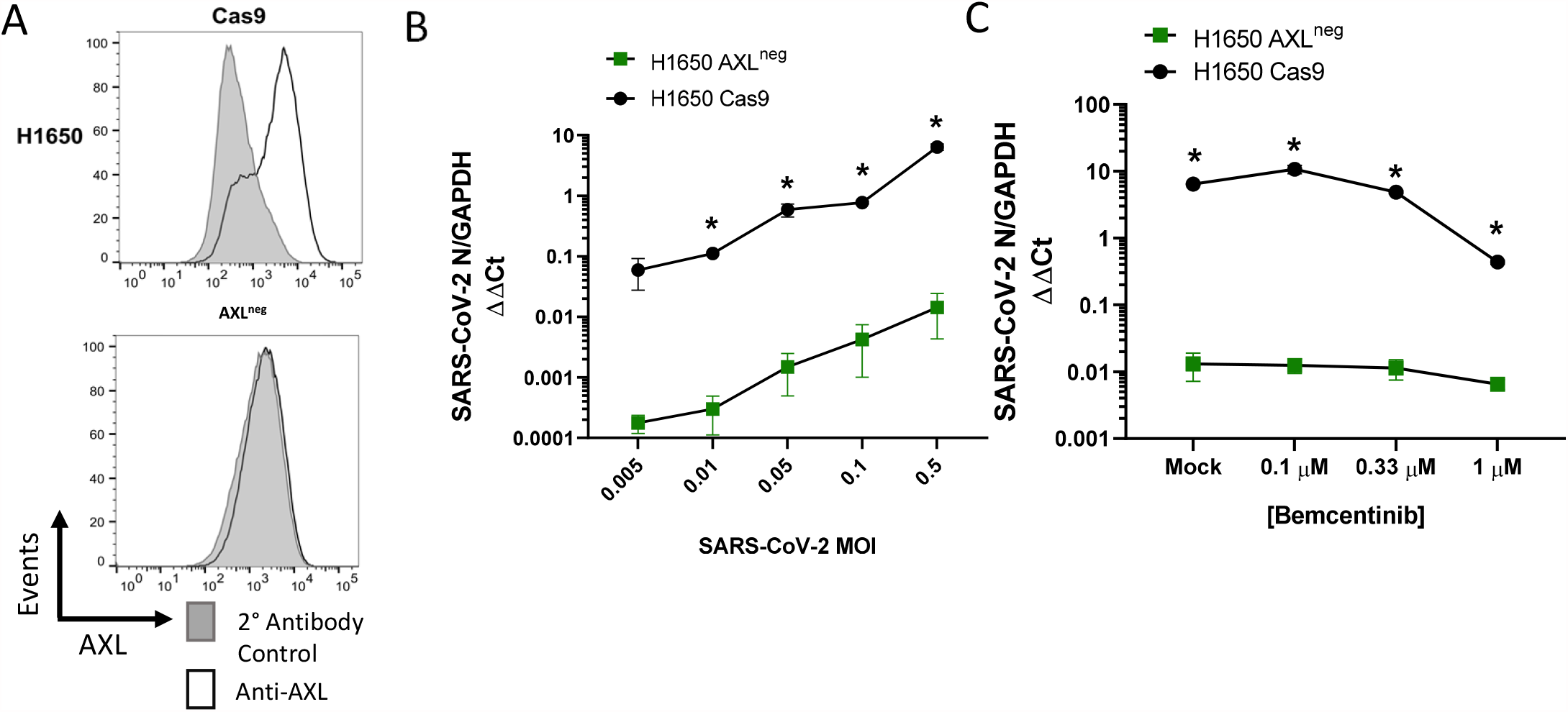
AXL knockout reduces viral loads and ablates inhibition by bemcentinib. **A)** H1650 AXL knockout cells were generated by lentiviral transduction of Cas9 and gRNA targeting AXL, followed by selection, enrichment, and biological cloning. Shown are flow cytometry histograms depicting AXL surface staining (black) and secondary only background (grey), demonstrating loss of AXL expression. **B)** H1650 AXL ^neg^ and H1650 Cas9 (parental) lines were challenged with SARS-CoV-2 at indicated MOIs. At 24hpi, viral loads were assessed by RT-qPCR. **C)** H1650 parental and AXL ^neg^ lines were treated with indicated concentration of bemcentinib for 1 hour and subsequently challenged with SARS-CoV-2 (MOI = 0.5) and viral loads determine by RT-qPCR 24 hpi. Data are pooled from at least 3 independent experiments (**B, C**) or are representative of at least 3 experiments (**A**). Data represented as means ± SEM. Multiple t-tests; asterisks represent p < 0.05.

### AXL facilitates infection of other betacoronaviruses

To assess the role of AXL in the related betacoronavirus, mouse hepatitis virus (MHV strain A59), we investigated the ability of bemcentinib to inhibit infection in C57BL/6J mouse bone marrow derived macrophages (BMDMs) that express AXL (33-35). As MHV uses the mouse receptor CEACAM as its cognate receptor, this model allowed us to examine the role of AXL with a coronavirus that is ACE2-independent (36). Bemcentinib added to BMDM cultures decreased virus load at 24 hours in a dose-dependent and an MOI-dependent manner (**S7A-B**). At higher MOIs the effect of bemcentinib was diminished. Bemcentinib treatment also inhibited MHV infection in peritoneal macrophages, an MHV permissive population phenotypically and functionally distinct from BMDMs (**S7C**). These studies provide evidence that AXL facilitates infection of multiple members of this enveloped virus family, independent of the cognate receptor used by the virus.

## DISCUSSION

Here, we demonstrate that PS receptors, AXL, TIM-1 and TIM-4, synergize with ACE2 to mediate SARS-CoV-2 infection of HEK 293T cells when ACE2, the cognate receptor for the virus, was expressed at low levels. PS receptors enhanced virion binding to cells in a PS-dependent manner. At higher levels of ACE2 expression, a role for the PS receptors was no longer observed. Similar findings were observed for TMPRSS2-facilitated, ACE2-dependent infection, indicating that when ACE2 is expressed on the plasma membrane at high concentrations these host proteins that assist SARS-CoV-2 entry are no longer required. A recent study reported that AXL mediates SARS-CoV-2 infection (19). That report suggested that AXL-mediated virus entry is independent of ACE2 and that AXL binds to the N-terminal domain of SARS-CoV-2 spike. Conclusions from our studies indicate that AXL and other PS receptors mediate enhancement of SARS-CoV-2 infection through interactions with virion-associated PS in an ACE2-dependent manner. We report several lines of evidence that are consistent with our contention that the PS receptors interact with virion-associated PS. First, PS liposomes abrogate binding and entry in a dose dependent manner. Second, interfering with PS/PS receptor complexes by mutating the TIM-1 PS binding pocket abrogates SARS-CoV-2 infection. These data are consistent with and support the well-established mechanism of PS receptor enhancement of enveloped virus infection (11-13, 37, 38). Third, we directly tested whether AXL binds to purified Spike or NTD by flow cytometry and biolayer interferometry assays and were unable to detect any interaction. Finally, the ability of diverse PS receptors to enhance SARS-CoV-2 infection in HEK 293T cells lend support to the contention that these receptors interact with PS rather than viral spike on the surface of SARS-CoV-2 to mediate productive infection. Thus, we conclude that AXL does not interact with SARS-CoV-2 spike, nor does it mediate virus entry unilaterally. It should be noted that this is the first example of an enveloped virus that utilizes PS receptors in conjunction with low/moderate expression of a high affinity surface receptor.

A previous study concluded that the related coronavirus, SARS-CoV, was not productively internalized by PS receptors (11). However, with insights from our studies, an alternative explanation is that PS receptor enhancement of coronavirus entry is ACE2-dependent and sufficient quantities of ACE2 on the plasma membrane abrogate a role for PS receptors. Thus, PS receptors only facilitate SARS-CoV-2 entry under conditions where ACE2 is expressed at suboptimal levels, conditions that were likely not evaluated in the cited study but are found on ACE2 expressing cells such as Vero E6 cells and many patient derived lung cell lines. As ACE2 expression is low within the lung, such suboptimal conditions may be highly relevant during SARS-CoV-2 infection (39, 40).

PS receptors have previously been shown to interact with PS on the surface of other enveloped viruses such as filoviruses, alphaviruses and flaviviruses and mediate internalization into endosomes (11, 41). However, PS receptor-dependent entry is a mechanism that is functionally out-competed by high-affinity viral glycoprotein-host receptor interactions, such as that of Lassa virus with α-dystroglycan (16, 20). In the case of Lassa virus entry PS receptors seem to serve as a backup entry mechanism, as these receptors only mediate virus internalization when the high affinity surface receptor, α-dystroglycan, is not expressed.

Our studies provide evidence that PS receptors enhance SARS-CoV-2 binding to cells and mediate internalization into endosomes where cysteine proteases potentiate spike protein triggering and subsequent fusion events. Consistent with the utilization of this uptake pathway, the cysteine protease inhibitor E-64 effectively blocked ACE2 or ACE2/PS receptor entry in HEK 293T cells. This is also supported by our super resolution microscopy observations that AXL and ACE2 colocalize during SARS-CoV-2 infection. However, in HEK 293T, the route of virus entry changes upon expression of TMPRSS2; virion entry is no longer sensitive to E-64. As others have reported, these findings are consistent with TMPRSS2-dependent entry dominating as the route of entry when TMPRSS2 is expressed (24, 42, 43). We also investigated virus infection of a variety of lung lines that endogenously express TMPRSS2, AXL, and ACE2. While findings with Calu-3 cells were similar to that we observed in TMPRSS2-transfected HEK 293T cells, the other TMPRSS2^+^ lung lines, such as H1650 cells, remained sensitive to E-64 and the AXL signaling inhibitor, bemcentinib. Differences in the ability of TMPRSS2 expression to control the route of entry may be due to a fine balance of surface expression of the various receptors and should be explored in more detail.

Our data indicate that AXL serves as the most important PS receptor for SARS-CoV-2 infection of the TIM and TAM families and our studies with MHV implicated AXL in facilitating infection of additional coronaviruses. While AXL is abundant on lung epithelial cells, it is also present in many organs in the body, with the exclusion of neural tissues (32, 44). Thus, it is likely a role for AXL in SARS-CoV-2 infection is not only relevant to lung cell populations, but ACE2-expressing tissues suspected to be affected by COVID-19 such as the heart and kidneys (45, 46). We surmise that AXL-inhibiting therapeutics could function in tandem with other antivirals, protecting a number of organs from infection. Our data suggest that the efficacy of bemcentinib will persist as the virus evolves, inhibiting the VOCs Alpha and Beta effectively. By targeting host proteins such as AXL we dramatically reduce the potential selection for pathogen mutants that reduce or abolish antiviral activity. Given that currently utilized small molecule therapeutics such as remdesivir targeting viral proteins have shown limited efficacy and the benefits of antibody-rich convalescent plasma is minimal, AXL inhibition by small molecule inhibitors such as bemcentinib offers a novel route of attack to reduce SARS-CoV-2 entry and disease (47, 48). The preferential utilization of AXL rather than TIM-1 by SARS-CoV-2 in Vero E6 cells was unexpected. In our previously studies, other enveloped viruses that utilize PS receptors, such as filoviruses, use TIM-1 preferentially when both proteins are expressed (12, 20). Further, a recent study identified that the TIM-1 IgV domain that contains the PS binding pocket serves as an effective inhibitor of enveloped virus infection regardless of the PS receptor utilized for virus uptake (49), consistent with the good affinity the TIM-1 PS binding pocket has for PS (50). Nonetheless, when TIM-1 is not present in cells and AXL is the sole PS receptor expressed, AXL is used by filoviruses and flaviviruses (11, 13, 51-53). The subpar utilization of AXL reported for other viruses may be due to the requirement for the adaptor protein, Gas6, to also be present, expression patterns, or steric factors. As multiple proteins from a variety of different PS receptors families can mediate uptake of apoptotic bodies, it is no surprise that PS receptor interactions with viruses are likewise intricate. Further studies are needed to understand the preferential use of AXL by SARS-CoV-2.

Bemcentinib, an orally bioavailable small molecule inhibitor of AXL, is currently in Phase II trials for non-small cell lung cancer (NSCLC) and a variety solid and hematological cancers (ClinicalTrials.gov IDs: NCT03184571, NCT03184558). However, multiple screens have identified bemcentinib as inhibitory to SARS-CoV-2 infection, bolstering this mechanism of entry (54, 55). Two phase 2 clinicals trial evaluating efficacy of bemcentinib in hospitalized COVID-19 patients are ongoing, with the first recently reporting short-term efficacy results (https://clinicaltrials.gov/ct2/show/NCT04890509). In this exploratory, open-label study bemcentinib was added to standard-of care (SoC) therapy to hospitalized patients (56). Though the primary endpoints (time to improvement by 2 points on WHO ordinal scale or time to discharge) showed a marginal benefit of bemcentinib treatment, there was evidence of potentially meaningful clinical benefit in a key secondary endpoint which was avoidance of deterioration. These interim data are promising and support further clinical investigations of this AXL inhibitor for treating SARS-CoV-2.

The robust body of PS receptor research completed in the last decade and historical patterns of zoonotic events (Ebola virus, Zika virus, coronaviruses) suggest that future emergent viral pathogens are likely to utilize PS receptors to enhance entry and infection. The observations reported here that PS receptors are utilized by a novel pandemic coronavirus support this conclusion. This confluence of information provides insights into a new class of potential therapeutics to stem future outbreaks, namely drugs aimed at inhibiting PS receptor activity.

Further studies are required to determine the role of TIM and TAM use *in vivo*; however, our studies demonstrate a role of PS receptors in SARS-CoV-2 infection in relevant cell populations and further extend the importance of PS receptors in enveloped virus entry to coronaviruses.

## STAR METHODS

## RESOURCE AVAILABILITY

### Lead contact

Further information and request for resources and reagents should be directed to and will be fulfilled by the lead contact Wendy J. Maury.

### Materials availability

This study did not generate new reagents.

### EXPERIMENTAL MODEL AND SUBJECT DETAILS

#### Ethics statement

This study was conducted in strict accordance with the Animal Welfare Act and the recommendations in the Guide for the Care and Use of Laboratory Animals of the National Institutes of Health (University of Iowa (UI) Institutional Assurance Number: #A3021-01). All animal procedures were approved by the UI Institutional Animal Care and Use Committee (IACUC) which oversees the administration of the IACUC protocols and the study was performed in accordance with the IACUC guidelines (Protocol #8011280, Filovirus glycoprotein/cellular protein interactions).

#### Mice

The mice (6-8 weeks old, female) used in these studies were obtained from the Jackson Laboratory (C57BL6/J). The protocol (#8011280) was approved by the Institutional Animal Care and Use Committee at the University of Iowa.

#### Primary Cells and Immortal Cell Lines

Bone marrow derived macrophages (BMDM) were isolated and cultured in RPMI-1640 supplemented with 10% Fetal Bovine Serum (FBS), 0.5 μg/mL of penicillin and streptomycin (pen/strep) and 50 ng/mL murine M-CSF. Vero E6 cells (ATCC CRL-1586), Vero TMPRSS2, Vero E6 and HEK 293T (ATCC CRL-11268) were cultured in Dulbecco’s modified Eagle’s medium (DMEM, GIBCO, Grand Island, NY) supplemented with 5-10% FBS and 1% penicillin/streptomycin (GIBCO). Blasticidin (5 μg/mL) was added to media supporting Vero E6 TMPRSS2 cell growth. A549^ACE2^ cells were generated by transduction of A549 (ATCC CCL-185) with a codon-optimized ACE2 encoding lentivirus and selection with 10 μg/mL blasticidin. Clonal populations were isolated and ACE2 expression verified by western blot. H1650, HCC2302, HCC1944 and H1819 human lung lines were maintained in RPMI with 5-10% FBS and pen/strep. Cell lines were periodically tested for mycoplasma contamination (E-Myco kit, Bulldog Bio, Portsmouth, NH) and cured of contamination before use (Plasmocin, Invitrogen, San Diego CA). Cell lines were authenticated periodically by ATCC (A549 and HEK 293T) or the lab responsible for their generation (H1650, HCC2302, HCC1944, H1819, and Vero TMPRSS2).

AXL knockout HCC2302 and H1650 were generated by transduction of parental cells with a Cas9 encoding lentivirus (kind gift of Aloysius Klingelhutz, University of Iowa) and selection in 10 μg/mL blasticidin for 10 days. Then cells were transduced with an Invitrogen LentiArray CRISPR gRNA lentivirus targeting AXL, and subsequently selected in puromycin at 2μg/mL for 5 days. Cells were then lifted, stained for AXL, and sorted for AXL^low^ cells at the University of Iowa Flow Cytometry Core on a FACSAria Fusion (Becton, Dickinson and Company, Franklin Lakes, New Jersey). Bulk populations of AXL^low^ cells were used for experiments as noted, and Clone #4 was generated by sorting single AXL^low^ cells into a 96 well plate. AXL expression was verified by flow cytometry on a FACSVerse (Becton, Dickinson and Company).

#### Viruses

Studies used the 2019n-CoV/USA-WA-1/2020 strain of SARS-CoV-2 (Accession number: MT985325.1) which was propagated on Vero E6 cells. Briefly, Vero E6 cells were inoculated with an MOI of 0.001 in DMEM supplemented with 2% FCS and pen/strep. Media was removed and refreshed 24 hpi. When cells exhibited severe cytopathic effect, generally 72 hpi, cells were freeze-thawed once, transferred to a conical tube, centrifuged at 1000g for 10 minutes, and supernatants were filtered through a 0.45 μm filter. Virus was sequenced via Sanger sequencing periodically for furin cleavage site mutations (none were detected) and only low passage stocks were used.

MHV (A59) stocks were generously provided by Dr. Stanley Perlman. Viral stocks were generated on Vero E6 and the TCID_50_ was determined on HeLa-mCECAM1 cells by identification of cytopathic effect at 5 days.

Stocks of recombinant vesicular stomatitis virus that expressed SARS-CoV-2 Spike containing the D614G mutation and nano-luciferase (rVSV/Spike) (kind gift of Dr. Melinda Brindley, Univ. GA) were generated in either Vero E6 or Vero E6 TMPRSS2 cells. Cells were infected with a low MOI (∼0.005) of virus and input was removed after ∼12 h. Upon evidence of cytopathology, supernatants were collected over a three-day period, filtered through a 0.45 μm filter and frozen at -80°C until purified. Supernatants were thawed and centrifuged overnight at 7000 x g to concentrate the virus. The virus pellet was resuspended in endotoxin-free PBS and layered over a 20% sucrose/PBS cushion. Virus was pelleted through the cushion by centrifugation at 28,000 rpm in a SW60Ti rotor (Beckman). The virus pellet was resuspended in PBS and the TCID50 was determined on Vero E6 cells.

All viral titers were determined by a modified Spearman-Karber method as previously described and reported as infectious units (IU)/mL (57).

## METHOD DETAILS

### Inhibitors

Bemcentinib (BGB324, R428) was provided by BerGenBio ASA (Bergen, Norway) and dissolved in DMSO for *in-vitro* studies. BMS777607 (Millipore Sigma) was dissolved in DMSO. E-64 (Millipore Sigma) was dissolved in DMSO.

### RNA isolation and qRT PCR

Total RNA for PCR was extracted from cells or tissue using TRIzol (Invitrogen, Cat# 15596018) according to the manufacturer’s protocol. Total isolated RNA (1μg) was reverse transcribed to cDNA using the High-Capacity cDNA Reverse Transcription Kit (Applied Biosystems Cat# 4368814). The resulting cDNA was used for amplification of selected genes by real-time quantitative PCR using Power SYBR Green Master Mix (Applied Biosystems, Cat# 4368708). Data were collected on QuantStudio 3 and Ct values determined with the QuantStudio Data Analysis software (Applied Biosystems). Averages from duplicate wells for each gene were used to calculate relative abundance of transcripts relative to housekeeping genes (HPRT, GAPDH, mouse cyclophilin) and presented as 2^-ΔΔCT^.

### VSV/Spike pseudovirus production

The production of SARS-CoV-2-Spike vesicular stomatitis virus (VSV/Spike) pseudovirions has been described previously (Hoffmann et al., 2020). Briefly, HEK 293T cells were seeded in 10 cm tissue culture plates (CellTreat; Ref# 229692). After 48 hours cells were transiently transfected with a SARS-CoV-2-Spike pCG1 plasmid (a kind gift from Dr. Stefan Pohlman as described in Hoffmann et al., 2020) using a standard polyethyleneimine (PEI) protocol. For this transfection, one tube was prepared with 16 μg of plasmid diluted in 1.5 mL of OPTI-Mem (Gibco; Ref# 31985-070). The second tube was prepared with PEI (1mg/mL) diluted in 1.5 mL of OPTI-Mem at a concentration of 3 μl/1 μg of DNA transfected. The tubes were then combined, vortexed for 10-15 seconds and left to incubate at room temperature for 15 minutes. The mixture was then added dropwise to the HEK 293T cells and returned to incubator overnight. Twenty-four hours after transfection the cells were infected with a stock of replication insufficient VSV virions expressing firefly luciferase that were pseudotyped with Lassa virus glycoprotein on the viral membrane surface. The infection was incubated for ∼6 hours at 37°C, was removed from the cells, and fresh media was added. Viral collection took place at 24- and 48-hours post-infection. Media supernatants were removed from the flasks, briefly spun down to remove cellular debris (180 x g for 1 minute) and filtered through a 0.45 μm syringe-tip disk PVDF filter (CellTreat; Ref# 229745). Supernatants were then concentrated by a 16-hour centrifugation step at 5380 x g at 4°C. Pseudovirions were purified through a 20% sucrose cushion via ultracentrifugation at 28,000 rpm for two hours at 10°C in a Beckman Coulter SW60Ti rotor. Pseudovirus was then resuspended in 1x PBS and stored at -80°C. Pseudovirus was titered using end point dilution on Vero E6 cells. All infections of cells using SARS-CoV-2-S pseudotyped virions were conducted at a volume of virus that gave a relative light unit (RLU) of roughly 100,000 – 200,000 RLU.

### HEK 293T transfections and plasmids

All transfections were performed in HEK 293T cells, with a total plasmid concentration of 2 μg. Cells were seeded into 6-well tissue culture plate (Dot Scientific; Ref# 667106) at a density of 5 × 10^5^ cells/well. Forty-eight hours after cell seeding, cells were transfected with CMV-driven expression vectors of ACE2, TIM-1, TIM-4, AXL, TYRO3, MerTK and TMPRSS2 (see plasmid details below) with a standard PEI transfection protocol. For this transfection one tube was prepared with plasmid DNA and 150 mM NaCl at a concentration of 25 μl/1 μg of DNA transfected. A second tube was prepared with 150 mM NaCl at a concentration of 25 μl/1 μg of DNA transfection along with PEI (1 mg/mL) at a concentration of 3 μl/ 1 μg of DNA transfected. Tubes were combined, vortexed vigorously for 10-15 seconds and incubated at room temperature for 15 minutes. Mixtures were added dropwise to HEK 293T cells. For all experiments using ACE2 50 ng of ACE2 plasmid were transfected, unless otherwise noted. For PS receptors, 1000ng of plasmid was transfected unless otherwise noted. All transfections were brought up to 2 μg total transfected DNA with a PCD3.1 empty expression vector.

### ACE2 and PS receptor expression detection via flow cytometry

To detect cell surface expression of ACE and PS receptor on transfected HEK293Ts, WT VeroE6s, and AXL knock down/out H1605 and HCC2302 clones, cells were lifted using 1x Versene (GIBCO; Ref# 15040066) at 37°C for ∼15 minutes and placed in 5mL polystyrene round-bottom tubes (Falcon Ref# 352052). Cells were washed once with FACS buffer (1x Sterile PBS, 2% FBS, 0.01% sodium azide). Cells were incubated for 30 minutes on ice with primary antibodies diluted in FACS buffer against ACE or PS receptors. Primary antibodies were diluted to 0.75 μg/mL in FACS buffer prior to incubation. Specific primary antibodies used as follows: goat anti-ACE2 (R&D AF933), goat anti-AXL (R&D154), goat anti-TIM-1 (R&D 1750), goat anti-Tyro3 (R&D AF859), goat anti-TIM-4 (R&D 2929), goat anti-MerTK (R&D AF891), rabbit anti-TMPRSS2 (Abcam ab92323). Cells were washed once with FACS buffer. Cells were then incubated with secondary antibodies at a 1:1000 dilution in FACS buffer on ice for 30 minutes. Secondaries used were donkey anti-goat IgG (H+L) Alexa Fluor 647 (Jackson Immuno Research; Ref# 705-605-003) and donkey anti-rabbit IgG (H+L) Alexa Fluor 647 (Invitrogen; A32733). Flowcytometry was performed on a Becton Dickinson FACS Calibur and analyzed by Flow Jo.

To examine both the surface and intracellular expression of hACE2 on H1650, Calu3, HCC1944, HCC2302, A549^ACE2^ and Vero E6 cells we performed the following protocol. Briefly, cells were staining with Fixable Viability Dye eFluor 780 (eBiosciences), goat anti-human ACE2 (R&D AF933) followed by secondary were donkey anti-goat IgG (H+L) Alexa Fluor 647 (Jackson Immuno Research; Ref# 705-605-003) To measure the intracellular expression of hACE2, cells were surface stained with Fixable Viability Dye eFluor 780, fixed (PFA 4%), permeabilized (1X PBS + 0.5%Tween20) and stained intracellularly using goat anti-human ACE2 (R&D AF933) followed by secondary donkey anti-goat labeled with AF647. Unstained cells, cells plus viability dye and cells plus secondary antibody/viability dye were included as a control in every staining. Samples were measured on a FACSverse cytometer (BD Biosciences) and data were analyzed with Flowjo software (BD Biosciences).

### VSV/SARS-CoV-2-spike pseudovirion studies with inhibitors

Following the transfection of HEK 293T cells, cells were incubated for 24 hours. At that time cells were lifted with 0.25% Trypsin (GIBCO; Ref# 25200-056) and plated at a density of 2 × 10^4^ cells/well on opaque, flat-bottomed, 96-well plates (Falcon; Ref# 353296). Each transfection was plated into at least 3 wells to create experimental replicates. Cells were incubated for an additional 24 hours. At that time, cells were infected with VSV-luciferase/SARS-CoV-2 Spike. Cells were incubated for an additional 24 hours. For experiments done with inhibitors, cells were treated with the concentrations of inhibitors noted in the figure panel immediately prior to being infected with pseudotyped virions. After 24 h, virus-containing media was removed and replaced with 35 μl of 1x Passive lysis buffer (Promega; Ref# E194A). Plates underwent three freeze-thaw cycles consisting of freezing on dry-ice for 15 minutes followed by thawing at 37°C for 15 minutes. We followed the protocol for measuring firefly luciferase as reported previously (Johnson et al., 2017). For this method 100 μl of luciferin buffer (100 μl of luciferin buffer (15 mM MgSO_4_, 15mM KPO_4_ [pH7.8], 1 mM ATP, and 1mM dithiothreitol) and 50 μl of 1mM d-luciferin potassium salt (Syd Laboratories; Ref# MB000102-R70170)) were added to each well and luminescence was read via Synergy H1 Hybrid reader (BioTek Instruments). Relative luminescence units were read out. Results analyzed by normalizing values to mock transfection with no protease inhibitors.

### HEK 293T SARS-CoV-2 infection studies

Following PEI transfection with plasmids as described in the previous section, cells were lifted with 0.25% trypsin and plated into 48-well plates at a density of 6 × 10^4^ (Dot Scientific; Ref# 667148). In our BSL3 facility, transfected HEK 293T cells were infected at a MOI = 0.5 with SARS-CoV-2. Cells were incubated at 37°C or 24 hours and then treated with TRIzol RNA isolation reagent and removed from the BSL3 facility. RNA extraction and cDNA generation proceeded as described. qRT-PCR was conducted on the cDNA using SARS-CoV-2-Spike and GAPDH primers. Data analyzed using the ΔΔCt method as described above.

### Virion binding assays

For Vero E6 binding studies, cells were grown to confluence in 48 well plates. Media was replaced with DMEM supplemented with 2% FBS and the indicated compounds. Cells were incubated at 10°C (preventing internalization and entry) until equilibrated and SARS-CoV-2 was added at MOI 5. Plates were returned to 10°C for 1 hour. Media was then removed and cells were washed three times with cold DPBS (GIBCO, Cat# 14190144), removing any unbound virus. Then 0.05% Trypsin-EDTA (GIBCO, Cat# 25300054) was added to control wells for 5 minutes at 37°C, and washed. After the final wash, all media was removed and replaced with TRIzol (Invitrogen, Cat# NCC1701D). RNA was isolated and analyzed as described. Binding studies in HEK 293T cells were performed using rVSV-SARS-CoV-2 Spike virions (rVSV/Spike) and binding was performed at room temperature in IMMULON 2HB flat bottom plates (Thermo Scientific, Waltham, MA).

### RNA sequencing and analysis

Following indicated treatments and infections that were performed in triplicate or quadruplicate, Vero E6 and A549^ACE2^ cells in 6 well formats were homogenized using QIAShredder tubes and total RNA isolated using the RNEasy kit with DNase treatment (Qiagen). High quality RNA samples that were verified by a Bioanalyzer (Agilent) was quantified and used as input to generate mRNA-seq libraries for the Illumina platform. Paired-end reads were performed at the Paired-end sequencing reads were subject to alignment to suitable reference genomes: human GRCh38 (GCA_000001405.15 - A549 cells), green monkey (Chlsab1/GCA_000409795.2 - Vero E6 cells) and SARS-CoV-2 (MN985325 - both A549 and Vero E6 cells). Alignments to human and monkey genomes were performed using hisat2 v2.0.5, while to viral genome using bowtie2 v2.2.9. Aligned reads were counted using featureCounts from subread package v1.5.2. Counted reads were normalized in R, using DESeq2 v1.30.0 and subjected to statistical analysis. The statistical analysis included computation of median based fold changes, Student t-test p values and false discovery rate (multiple testing correction).

### Purified spike protein flow cytometry binding studies

The NTD-Fc and RBD-Fc constructs were kindly provided by Tom Gallagher. They contain the Fc region of human IgG1 fused to the N-terminal domain of SARS-CoV-2 Spike (residues 1-309) or the RBD-containing C-terminal domain of the S1 subunit (residues 310-529). We also generated an Fc-Spike construct that contains the Fc region of human IgG1 fused to a cleavage-negative form of the Spike ectodomain (subunits S1 and S2, corresponding to positions 1-1274). To eliminate the polybasic furin cleavage site of Spike, we replaced the Arg-Arg-Ala-Arg motif at positions 682-685 with Ser-Ser-Ala-Ser. All proteins were produced by transient transfection of 293T cells using polyethylenimine. Proteins were harvested in 293S ProCDM and purified using Protein A beads. Eluted products were dialyzed against phosphate buffered saline (pH 7.4). All proteins were analyzed by SDS-PAGE and gels were silver stained to verify the purity of the eluted product.

We measured the binding efficiency of anti-NTD antibody AM121 (Acro Biosystems) to the Spike-based constructs using an enzyme-linked immunosorbent assay (ELISA), as previously described (58, 59). For this purpose, the NTD-Fc, RBD-Fc or Spike-Fc suspended in PBS were attached to 96-well protein-binding plates by incubation at isomolar concentrations (2, 1.37 and 5 μg/mL of the probes, respectively). The next day, wells were washed once with buffer containing 140 mM NaCl, 1.8 mM CaCl2, 1 mM MgCl2, 25 mM Tris pH 7.5, 20 mg/mL BSA and 1.1% low-fat milk. The anti-NTD antibody suspended in the same buffer was then added to the wells at 0.5 μg/mL. Binding of the anti-NTD antibody was detected using a goat anti-human kappa light chain conjugated to horseradish peroxidase (HPR) (BioRad). To normalize for the amount of the bound probes, we also quantified the amount of probe bound to the wells by incubation with an HRP-conjugated goat anti-human antibody preparation. Binding of the HRP-conjugated antibodies was measured by luminescence using SuperSignal West Pico enhanced chemiluminescence reagents and a Synergy H1 microplate reader, as previously described (60).

To determine binding of the above probes to AXL, we used flow cytometry. Briefly, HEK 293T cells were seeded in 6-well plates (8.5E5 cells per well) and transfected the next day with 1.5 μg of empty vector or plasmids that express AXL or the full-length form of human angiotensin converting enzyme 2 (ACE2) using JetPrime transfection reagent (PolyPlus). Three days after transfection, cells were detached using PBS supplemented with 7.5 mM EDTA and washed once with washing buffer (PBS supplemented with 5% newborn calf serum). Cells were then incubated with the NTD-Fc, RBD-Fc or Spike-Fc probes (at 5 μg/mL) or anti-AXL antibody (at 0.75 μg/mL) in the same buffer for one hour on ice and were washed four times with washing buffer. To detect binding of the Fc probes, we used a goat anti-human polyclonal antibody preparation conjugated to Alexa 647. To detect binding of the anti-AXL antibody, we used a goat anti-donkey polyclonal antibody preparation conjugated to Alexa 594. Secondary antibodies were added at a 1:500 dilution and incubated with the cells on ice for one hour. Cells were then washed and analyzed by flow cytometry. Staining was evaluated on a Becton Dickinson FACS Calibur and analyzed by Flow Jo.

### Biolayer Interferometry

Biolayer interferometry was performed on an OctetRed96 (Pall Forte-Bio, USA) using NiNTA-(Forte-Bio 185101) or Streptavidin- (Forte-Bio 18-5019) coated Dip and Read biosensors for immobilisation of His-tagged or biotin-tagged proteins respectively. All samples were diluted in Kinetic Buffer (0.1% BSA, 0.02% Tween 20, 0.05% Sodium azide in PBS). Biosensors were equilibrated in Kinetic Buffer, and protein (His-tagged SARS-Cov-2 S1 protein NTD, Acro Biosystems, S1D-C52H6; irrelevant control His-tagged β4 integrin fibronectin type III domain, generous gift from Petri Kursala laboratory; biotinylated tilvestamab anti-AXL antibody, BerGenBio, Norway) at a concentration of 0.7μM was loaded for 10 minutes. A baseline was taken for 2 minutes in Kinetic Buffer before testing association of the target binding protein for 10 minutes (AXL extracellular domain fused to human IgG Fc region, BerGenBio ASA, 2μM; positive control Anti-SARS-Cov2 spike NTD Neutralizing antibody, Acro Biosystems, AM121, 0.1μM; IgG1 isotype control antibody, BioXcell, BE0297, 0.1μM). Results were analyzed in Prism 9.1.1 for MacOS (GraphPad Software, USA) by aligning to the baseline values immediately prior to association and subtracting signal from isotype control.

### STED sample preparation and image acquisition

12mm #1.5 coverslips (Fisher Scientific; Ref# 12-545-81P) were coated with 0.1% bovine Achilles’ tendon collagen diluted in 1x sterile PBS. Collaged solution was incubated at 37°C for 2 hours before being plated to 12mm coverslips. Collagen was allowed to incubate on coverslips 12 hours at 37°C. Slips were then rinsed with 1x PBS and dried in 37°C incubator. Slips were stored in sterile 1x PBS until use. Vero E6 cells were plated onto collaged coated slips at 30,000 cells/slip. 24 hours after plating, cells were either left uninfected, or infected with SARS-CoV02 (WA-1) at an MOI = 0.01 and incubated for 24 hours. At that time cells were washed once with sterile 1x PBS and then fixed with 4% PFA solution (Electron Microscopy Sciences; Ref# 15710) for 10 minutes at room temperature. Following PFA fixation, cells were washed three times with 1x PBS and stored at 4°C until use.

For immunofluorescent staining, coverslips were incubated for 2 hours at RT with a blocking buffer consisting of 1% Triton X-100, 0.5% sodium deoxycholate, 1% egg albumin and 0.01% sodium azide all suspended in 1x PBS. After 2-hour blockade, coverslips were incubated overnight at 4°C with primary antibodies against hACE2 (goat anti-hACE2; R&D AF933), and hAXL (rabbit anti-hAXL; Cell Signaling C89E7). Cells were washed three times with 1x PBS for 5 minutes each. Samples with goat anti-hACE2 primary antibodies were first incubated with donkey anti-goat IgG (H+L) Alexa Fluor 568 (Invitrogen; A-11057) for 1 hour at RT. Cells were washed three times with 1x PBS for 5 minutes. Coverslips incubated with donkey anti-goat Alexa Fluor 568 were then incubated with 5% NGS (Sigma; G9023) for one hour at RT. Cells were washed three times 1x PBS then incubated for one-hour room temperature with goat anti-rabbit IgG (H+L) Alexa Fluor Plus 488 (Invitrogen; A32731TR). After washing three times for 5 minutes with 1x PBS cells were fixed to glass microscopy slides (Fisher Scientific; 12-550-15) with 12ul of Prolong Glass mounting medium (Invitrogen; P36982). Coverslips cured for 48 hours before imaging.

Image acquisition was performed on the Leica SP8 3X STED confocal microscope equipped with an HC PL APO C32 100X/1.4 oil objective lens, and LAS X software (Leica Microsystems; version 3.5.5.19976) in the Central Microscopy Research Facility at the University of Iowa. Excitation was performed using a white light laser set to 20% intensity. Depletion was performed with a 660nm laser set at 7% intensity for Alexa Fluor 488, and 775 nm with 16.5% laser for Alexa Fluor 568 fluorophore. Depletion lasers were aligned using the auto STED alignment tool in the LAS X software. Laser strength and gain were adjusted to prevent pixel saturation. Images collected with two-line averages and 2 frame accumulations. Post image analyses include deconvolution using Huygens Professional Software and the deconvolution Wizard auto functionality (Scientific Volume Imaging). Fluorescence intensity plot profiles were created using ColorProfiler plug-in for ImageJ (Dimiter Prodanov). Pearson’s correlation coefficients were performed by using freehand ROI selection tool in ImageJ to outline individual cells and performing colocalization calculations using the Colocalization Test plugin (Tony Collins et al.). Depicted are the R values for 20 cells across five separate fields imaged from one coverslip.

## QUANTIFICATION AND STATISTICAL ANALYSIS

Statistical analysis was completed in GraphPad Prism v9.0.2 (GraphPad Software, San Diego, CA). Quantification of flow cytometry data was completed in FlowJo v10.7.1 (Becton, Dickinson & Company, Ashland, OR). Statistical significance was defined as p < 0.05, and denoted by a single asterisk (*). Details regarding statistical tests used and exact values of n can be found in the corresponding figure legends. All data presented is representative of n = 3 independent experiments unless otherwise noted.

## AUTHOR CONTRIBUTIONS

See Supplemental Figure 8 for author contributions matrix.

## ACKNOWLEDGEMENTS

This study was primarily supported by NIH/NIAID grant R01 AI134733 (WJM), NIH/NCI grants P50 CA070907 (JM), U54 CA260560 (JM) and a contract from BerGenBio to WJM. DB was supported by NIH grant T32AI007511 (Training in Mechanisms of Parasitism). HVE was supported by T32 GM007337.

Kaitlyn Bohan graciously designed and created our graphical abstract image. The Cas9/BlastR lentiviral expression vector was generously provided by Aloysius Klingelhutz, PhD. The A549^ACE2^ cell line was kindly provided by Balaji Manicassamy, PhD. Paige Richards assisted isolation of the H1650 AXL^neg^ clone.

Declaration of Interests: G.G., D.M., and E.C. are employees of BerGenBio ASA, a company with financial interests in this field. Partial funding was provided by BerGenBio ASA. No other authors have competing interests to declare. The funders had no role in study design, data collection, data analysis, decision to publish, or preparation of the manuscript.

Data presented herein were obtained at the Genomics Division of the Iowa Institute of Human Genetics which is supported, in part, by the University of Iowa Carver College of Medicine. The authors would like to acknowledge vital assistance from Chantal Allamargot, PhD. Imaging was performed using the Leica SP8 STED super resolution confocal available for use in the microscopy core.

The data presented herein were obtained at the Flow Cytometry Facility, which is a Carver College of Medicine / Holden Comprehensive Cancer Center core research facility at the University of Iowa. The facility is funded through user fees and the generous financial support of the Carver College of Medicine, Holden Comprehensive Cancer Center, and Iowa City Veteran’s Administration Medical Center.

## SUPPLEMENTAL FIGURE LEGENDS

**Supplemental Figure 1: PS receptors synergize with ACE2, enhancing SARS-CoV-2 infection of HEK 293T cells. A)** Representative surface staining of receptors transfected into cells. **B)** Surface expression (MFI) of proteins in mock transfected (empty vector) and transfected HEK 293T at 48 hours after transfection. Background fluorescence is shown for secondary antibodies used in experiment (α-goat or rabbit secondaries). **C)** HEK 293T cells, transfected PS receptors as noted with or without 250 ng of ACE2 were transduced with rVSV-luciferase/Spike. Transduction was assessed 24 hours later via luminescence. **D)** Expression of MerTK did not affect rVSV-luciferase/Spike transduction in the presence of 250 ng of transfected ACE2 plasmid. **E)** Expression of ACE2, TIM-1 or AXL did not enhance infection of VSV-luciferase/Lassa virus GP pseudovirions. HEK 293T cells were transfected with PS receptor plasmids and 50 ng of ACE2 and infected 48 hours later. Panels C-E are shown as fold change of luciferase activity in cell lysates relative to mock transfected lysates that were set to a value of 1.

Data shown are pooled from at least three independent experiments (**C, D**, and **E)**. Data represented as means ± SEM. One-Way ANOVA with multiple comparisons (**C, D**), Student’s t-test (**E**); asterisks represent p < 0.05.

**Supplemental Figure 2: PS receptors interact with SARS-CoV-2 by binding to PS. A**) AXL surface expression in transfected HEK 293T cells. **B**) Soluble purified S1/S2-Fc and NTD-Fc are detected by an NTD monoclonal antibody by ELISA. **C**) All spike-Fc proteins bind and are detected at equivalent levels of ELISA plates.

**Supplemental Figure 3: The route of SARS-CoV-2 entry is altered by TMPRSS2 expression**. ATPLite cytotoxicity assay in H1650 cells, 24 hours following treatment with E64. Data are represented as means +/-SEM.

**Supplemental Figure 4: AXL has a prominent role in SARS-CoV-2 entry in Vero E6 cells.A)** ACE2, AXL and TIM-1 surface expression MFI in Vero E6 cells, as assessed by flow cytometry. Background fluorescence is shown for secondary antibodies used in experiment. **B)** Cell surface versus intracellular ACE2 expression in VeroE6 cells. Indicated cells were lifted, permeabilized as noted, and stained with anti-ACE2 unconjugated primary antibodies and Alexa 647 secondaries. **C)** Bemcentinib toxicity 24 hours after treatment was measured by ATPlite assay in H1650 cell line. **D)** VSV-GFP/Spike entry was measured by flow cytometry 24 hours after challenge to Vero E6 cells treated with bemcentinib. **E)** Vero E6 were treated with ARD5 (TIM-1 blocking antibody) 1 hour before infection with rVSV /SARS-CoV-2 Spike or rVSV /EBOV-GP (MOI = 0.01). Viral load was measured 24 hpi by RT-qPCR. **F)** Plot profiles of ACE2 and AXL intensity are shown in from STED micrographs in Fig. **4G**, representing signal intensity along the yellow lines in the merged panels.

Data in A, C, D and E are shown as means ± SEM. Multiple t-tests were performed in C and Student’s t-test was performed in D and E; asterisks represent p < 0.05.

**Supplemental Figure 5: AXL inhibition reduces SARS-CoV-2 infection in human lung cells. A)** Multiple SARS-CoV-2 permissive cell lines were stained for extracellular ACE2, AXL, TIM-1, and TMPRSS2 protein, and expression was quantified by flow cytometry. Shown are flow cytometry histograms depicting target surface staining (black line) and secondary only background (grey shade) **B)** H1650 cells were treated with 1 μM of bemcentinib and infected with one of three different variants of SARS-CoV-2: WA-1; B.1.1.7 or B.1.351 (MOI=0.5 for all variants). RNA was isolated at 24 hpi and assessed for virus load. **C)** H1650 cells were infected with SARS-CoV-2 (MOI = 0.5) after treatment with the indicated concentration of camostat for 1 hour. Viral loads 24hpi were measured by qRT-PCR. **D)** Extracellular and intracellular staining of ACE2 are shown in multiple cell lines. Presented as frequency positive cells.

Data represented as means ± SEM. Student’s t-test; asterisks represent p < 0.05. **Supplemental Figure 6: AXL knockout reduces viral loads and ablates inhibition by bemcentinib. A)** H1650 and HCC2302 AXL knockout cells were generated by lentiviral transduction of Cas9 and gRNA targeting AXL, followed by selection. These are designated “Bulk AXL^low^” Shown are flow cytometry histograms depicting AXL surface staining (black) and secondary only background (grey), demonstrating complete loss of AXL expression in H1650 AXL^neg^. **B)** H1650 AXL^low^ and H1650 Cas9 (parental) lines were challenged with SARS-CoV-2 at indicated MOIs for 24 hpi and viral loads assessed by RT-qPCR. **C)** H1650 parental and AXL^low^ lines were treated with indicated concentration of bemcentinib for 1 hour and subsequently challenged with SARS-CoV-2 (MOI = 0.5) and viral loads determine by RT-qPCR 24 hpi. **D-E)** As in B-C with HCC2302 cells.

Data are pooled from at least 3 independent experiments (**B, C, D, E**) or are representative of at least 3 experiments (**A**). Data represented as means ± SEM. Multiple t-tests were performed; asterisks represent p < 0.05.

**Supplemental Figure 7: A)** Bone marrow derived macrophages from C57bl6/J mice were treated as indicated for 1 hour, challenged with MHV (stain A59) at the indicated MOI. Viral loads were assessed 24 hpi by RT-qPCR. **B)** BMDMs were treated with indicated concentrations of bemcentinib for 1 hour, infected with MHV (MOI = 0.001) for 24 hours and viral load assessed by RT-qPCR. **C)** As in B, MHV infection of peritoneal macrophages (MOI =0.001) treated with indicated concentrations of bemcentinib.

Data shown are representative of 3 independent experiments. Data represented as means ± SEM. Student’s t-test; asterisks represent p < 0.05.

