## Supplemental Figures for "Phosphatidylserine Receptors Enhance SARS-CoV-2 Infection: AXL as a Therapeutic Target for COVID-19"

S1: Supplemental data related to Figure 1

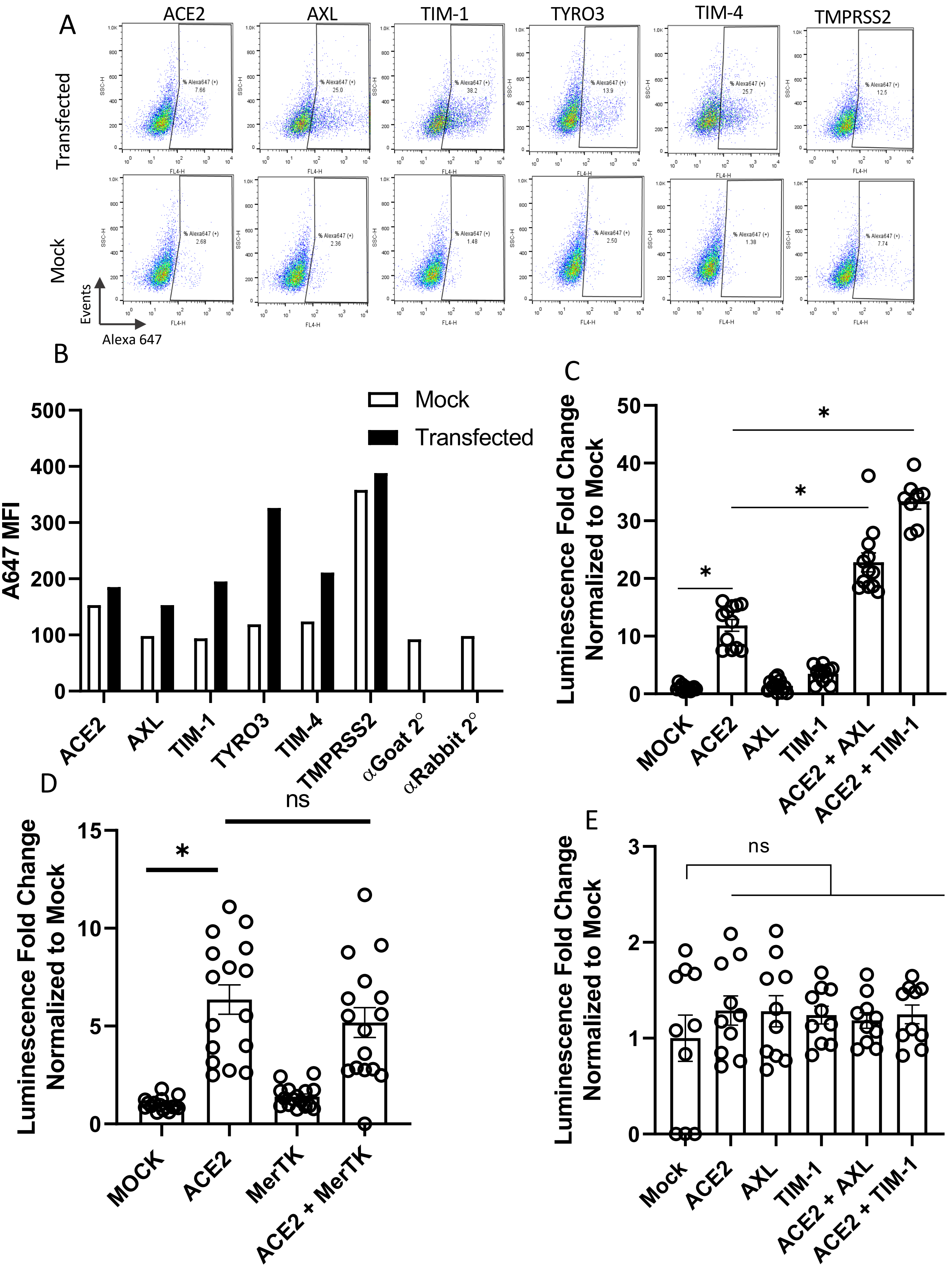

**Supplemental Figure 1: PS receptors synergize with ACE2, enhancing SARS-CoV-2 infection of HEK 293T cells**. **A)** Representative surface staining of receptors transfected into cells. **B)** Surface expression (MFI) of proteins in mock transfected (empty vector) and transfected HEK 293T at 48 hours after transfection**.** Background fluorescence is shown for secondary antibodies used in experiment (α-goat or rabbit secondaries). **C)** HEK 293T cells, transfected PS receptors as noted with or without 250 ng of ACE2 were transduced with rVSV-luciferase/Spike. Transduction was assessed 24 hours later via luminescence. **D)** Expression of MerTK did not affect rVSV-luciferase/Spike transduction in the presence of 250 ng of transfected ACE2 plasmid. **E)** Expression of ACE2, TIM-1 or AXL did not enhance infection of VSV-luciferase/Lassa virus GP pseudovirions. HEK 293T cells were transfected with PS receptor plasmids and 50 ng of ACE2 and infected 48 hours later**.** Panels C-E are shown as fold change of luciferase activity in cell lysates relative to mock transfected lysates that were set to a value of 1.

Data shown are pooled from at least three independent experiments (**C**, **D**, and **E)**. Data represented as means ± SEM. One-Way ANOVA with multiple comparisons (**C, D**), Student's t-test (**E**); asterisks represent $p < 0.05$.

S2: Supplemental data related to Figure 2

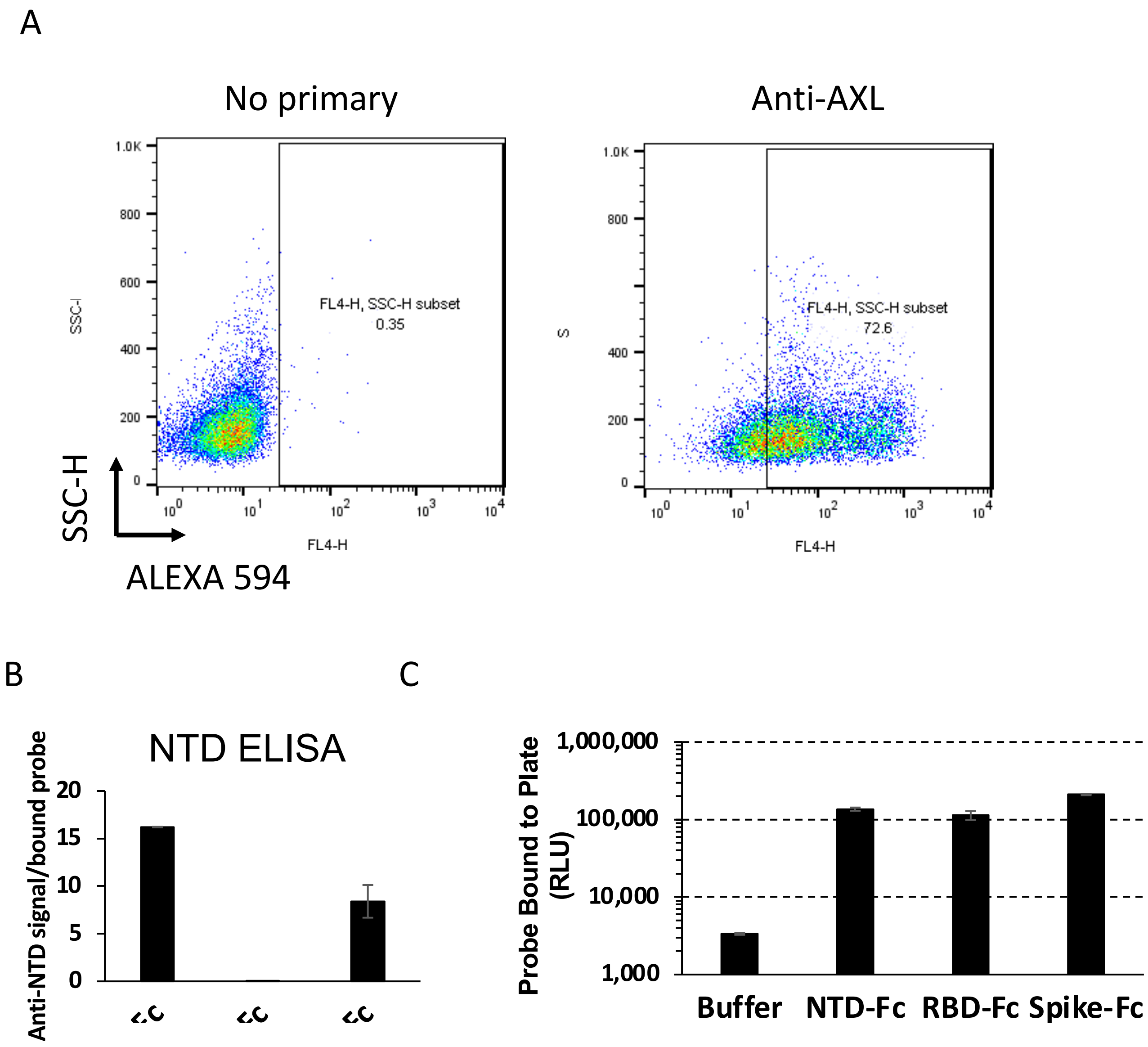


**Supplemental Figure 2: PS receptors interact with SARS-CoV-2 by binding to PS.** **A**) AXL surface expression in transfected HEK 293T cells. **B**) Soluble purified S1/S2-Fc and NTD-Fc are detected by an NTD monoclonal antibody by ELISA. **C**) All spike-Fc proteins bind and are detected at equivalent levels of ELISA plates.

S3: Supplemental data associated with Figure 3

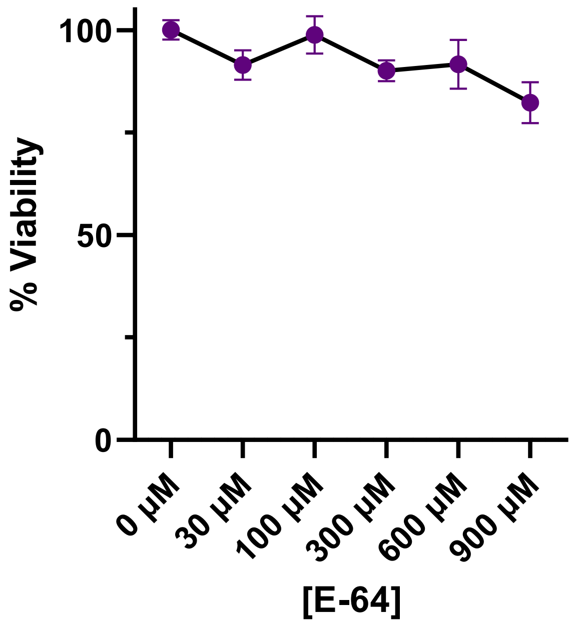


**Supplemental Figure 3: The route of SARS-CoV-2 entry is altered by TMPRSS2 expression.** ATPLite cytotoxicity assay in H1650 cells, 24 hours following treatment with E64. Data are represented as means +/- SEM.

S4: Supplemental data associated with Figure 4

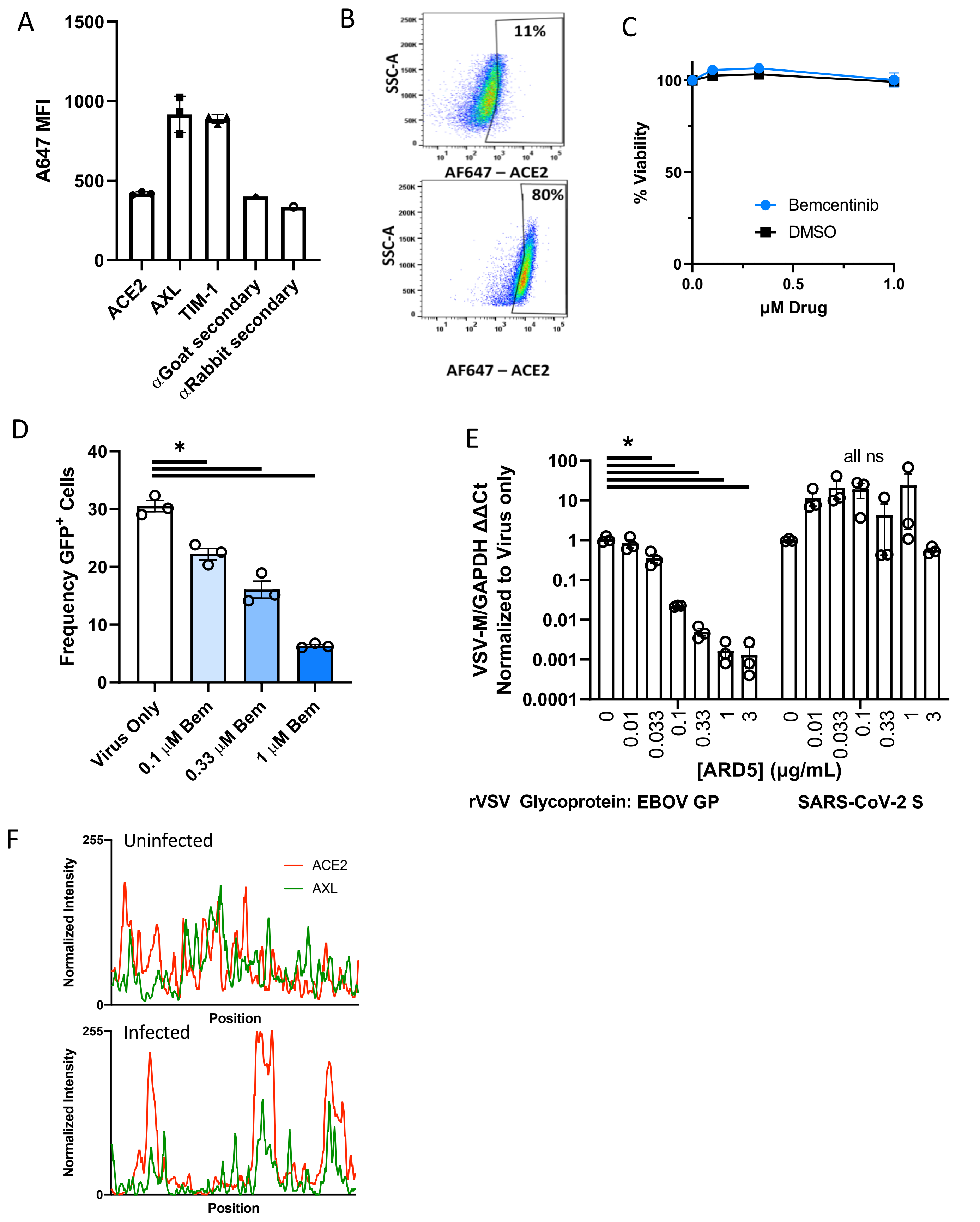

**Supplemental Figure 4: AXL has a prominent role in SARS-CoV-2 entry in Vero E6 cells**. **A)** ACE2, AXL and TIM-1 surface expression MFI in Vero E6 cells, as assessed by flow cytometry. Background fluorescence is shown for secondary antibodies used in experiment. **B)** Cell surface versus intracellular ACE2 expression in VeroE6 cells. Indicated cells were lifted, permeabilized as noted, and stained with anti-ACE2 unconjugated primary antibodies and Alexa 647 secondaries. **C)** Bemcentinib toxicity 24 hours after treatment was measured by ATPlite assay in H1650 cell line. **D)** VSV-GFP/Spike entry was measured by flow cytometry 24 hours after challenge to Vero E6 cells treated with bemcentinib. **E)** Vero E6 were treated with ARD5 (TIM-1 blocking antibody) 1 hour before infection with rVSV /SARS-CoV-2 Spike or rVSV /EBOV-GP (MOI = 0.01). Viral load was measured 24 hpi by RT-qPCR. **F)** Plot profiles of ACE2 and AXL intensity are shown in from STED micrographs in Fig. **4G**, representing signal intensity along the yellow lines in the merged panels.

Data in A, C, D and E are shown as means ± SEM. Multiple t-tests were performed in C and Student's t-test was performed in D and E; asterisks represent $p < 0.05$.

.

### S5: Supplemental data related to Figure 5

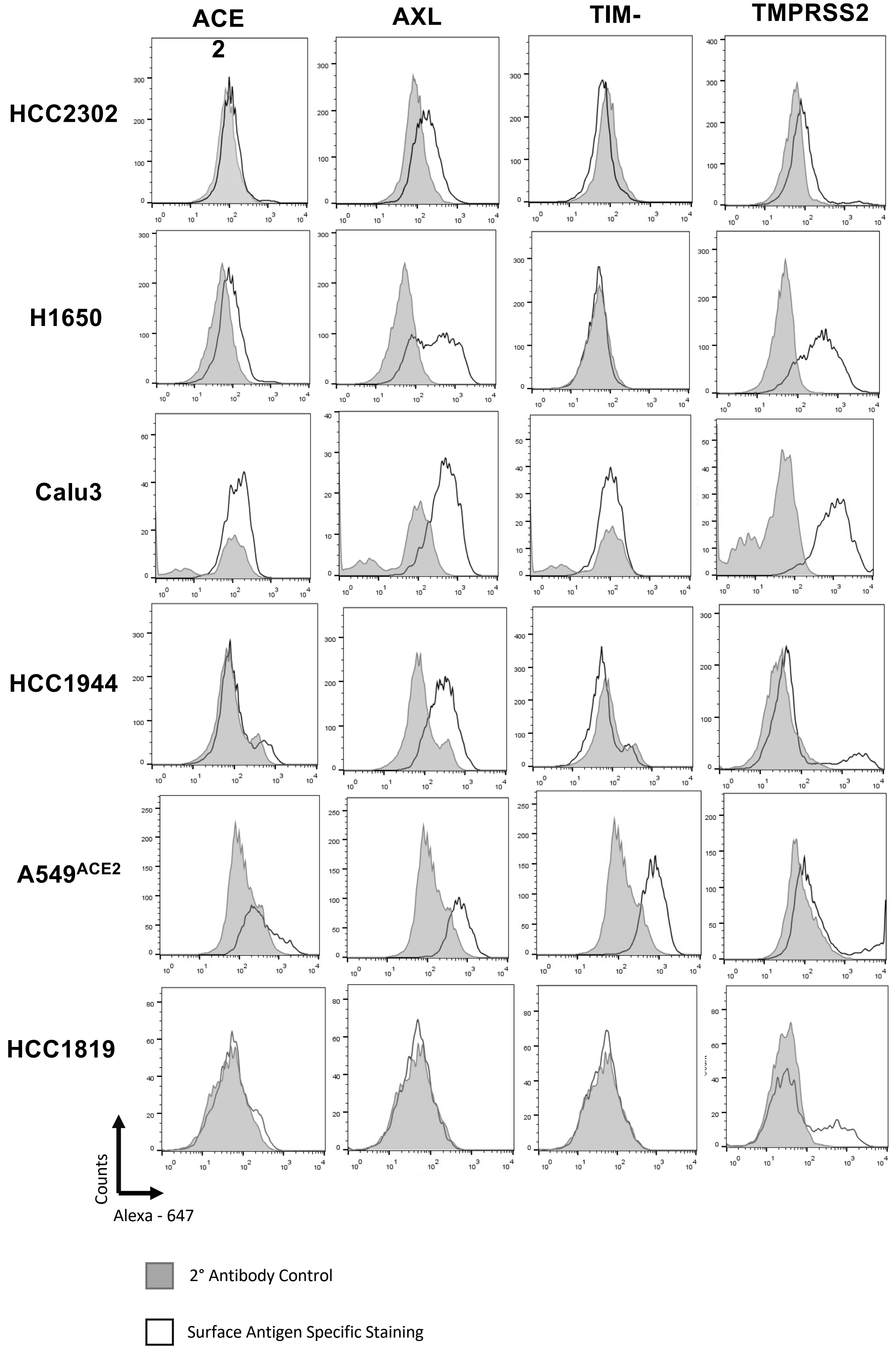

S5 con't: Supplemental data related to Figure 5

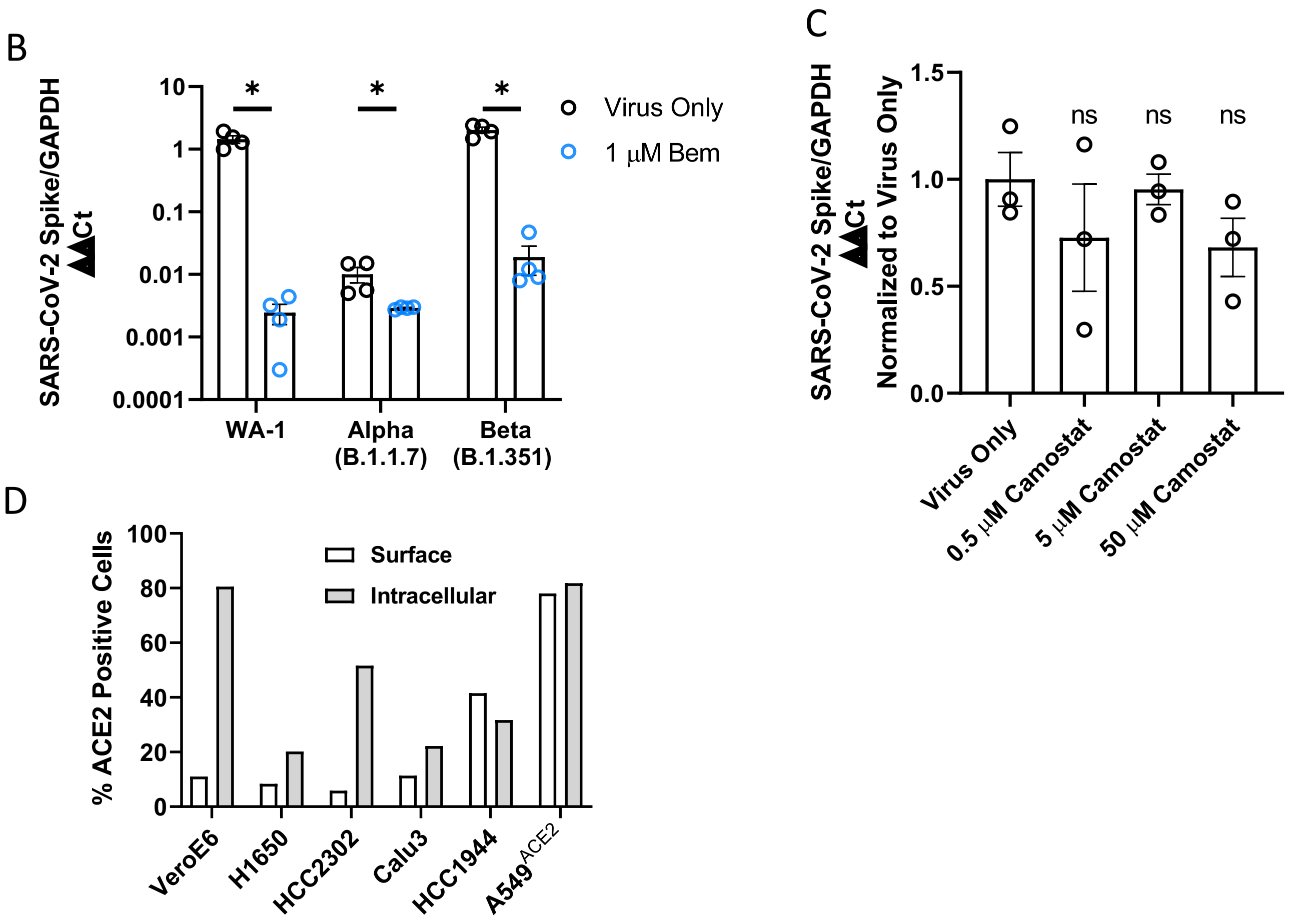


**Supplemental Figure 5: AXL inhibition reduces SARS-CoV-2 infection in human lung cells. A)** Multiple SARS-CoV-2 permissive cell lines were stained for extracellular ACE2, AXL, TIM-1, and TMPRSS2 protein, and expression was quantified by flow cytometry. Shown are flow cytometry histograms depicting target surface staining (black line) and secondary only background (grey shade) **B)** H1650 cells were treated with 1 µM of bemcentinib and infected with one of three different variants of SARS-CoV-2: WA-1; B.1.1.7 or B.1.351 (MOI=0.5 for all variants). RNA was isolated at 24 hpi and assessed for virus load. **C)** H1650 cells were infected with SARS-CoV-2 (MOI = 0.5) after treatment with the indicated concentration of camostat for 1 hour. Viral loads 24hpi were measured by qRT-PCR. **D)** Extracellular and intracellular staining of ACE2 are shown in multiple cell lines. Presented as frequency positive cells.
Data represented as means ± SEM. Student's t-test; asterisks represent $p < 0.05$.

### S6: Supplemental data related to Figure 6

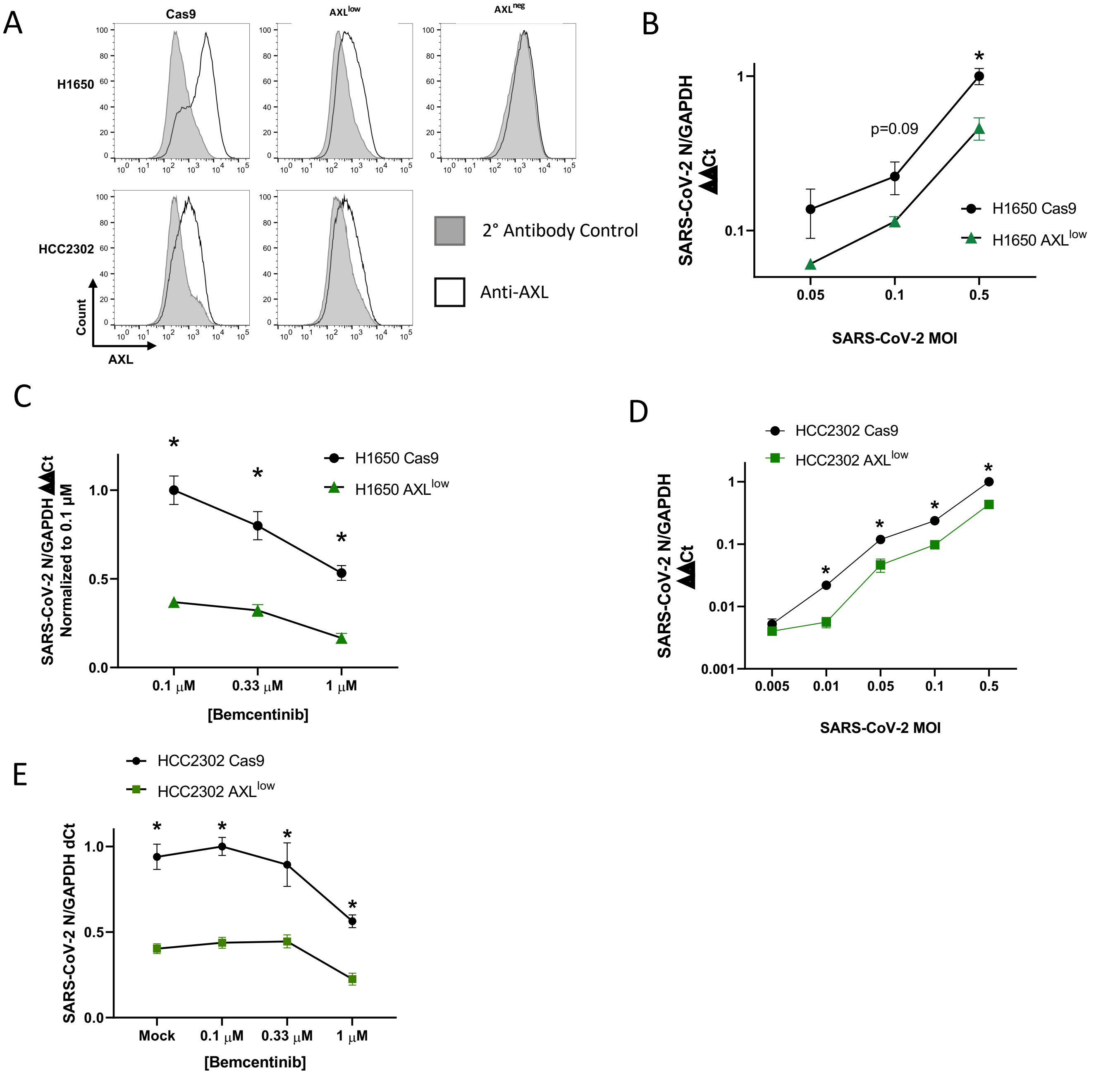

**Supplemental Figure 6: AXL knockout reduces viral loads and ablates inhibition by bemcentinib. A)** H1650 and HCC2302 AXL knockout cells were generated by lentiviral transduction of Cas9 and gRNA targeting AXL, followed by selection. These are designated “Bulk $AXL^{low}$” Shown are flow cytometry histograms depicting AXL surface staining (black) and secondary only background (grey), demonstrating complete loss of AXL expression in H1650 $AXL^{neg}$. **B)** H1650 $AXL^{low}$ and H1650 Cas9 (parental) lines were challenged with SARS-CoV-2 at indicated MOIs for 24 hpi and viral loads assessed by RT-qPCR. **C)** H1650 parental and $AXL^{low}$ lines were treated with indicated concentration of bemcentinib for 1 hour and subsequently challenged with SARS-CoV-2 (MOI = 0.5) and viral loads determine by RT-qPCR 24 hpi. **D-E)** As in B-C with HCC2302 cells.

Data are pooled from at least 3 independent experiments (**B**, **C**, **D**, **E**) or are representative of at least 3 experiments (**A**). Data represented as means ± SEM. Multiple t-tests were performed;  asterisks represent $p < 0.05$.

S7: Mouse Hepatitis Virus

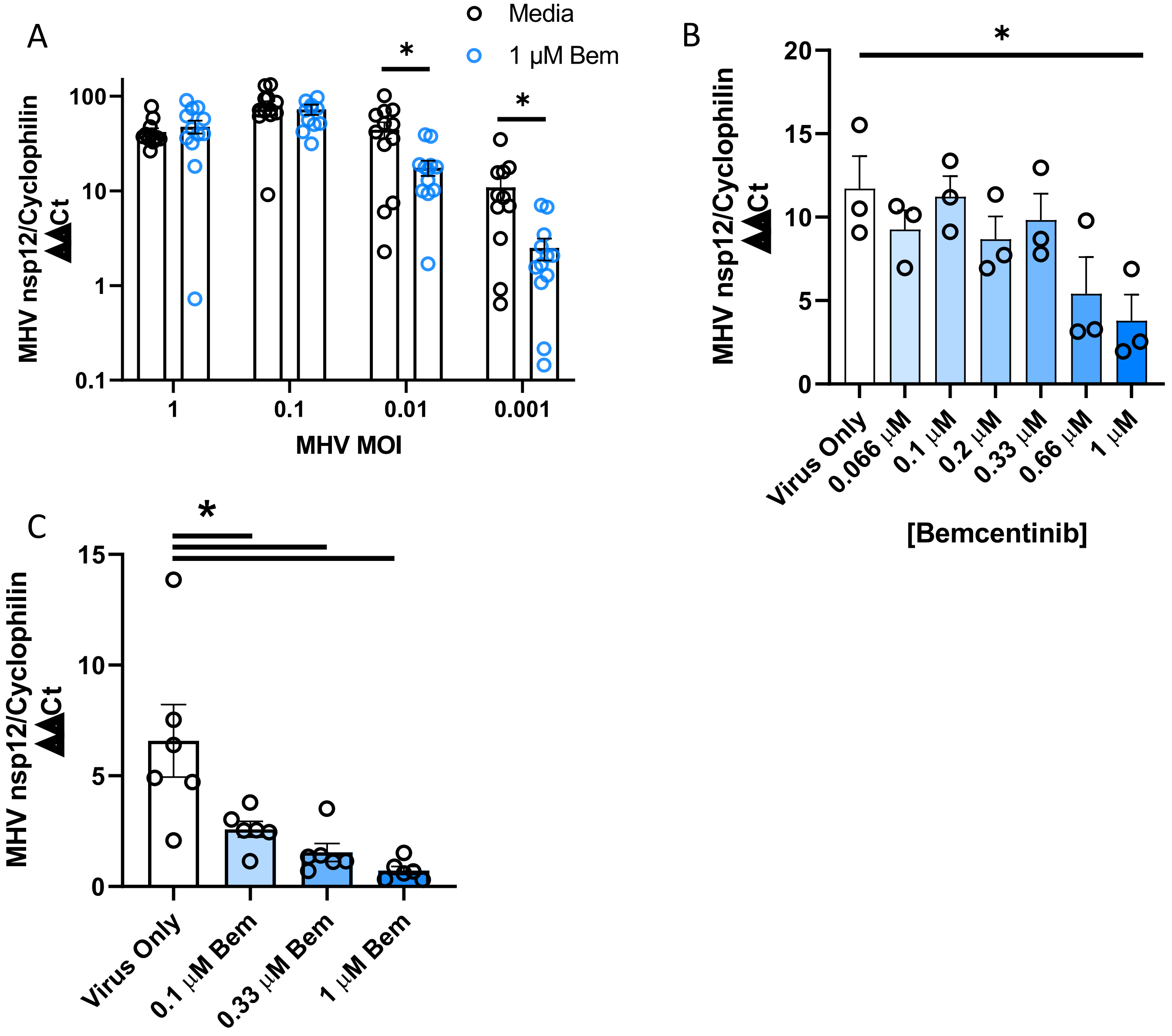


**Supplemental Figure 7: A)** Bone marrow derived macrophages from C57bl6/J mice were treated as indicated for 1 hour, challenged with MHV (stain A59) at the indicated MOI. Viral loads were assessed 24 hpi by RT-qPCR. **B)** BMDMs were treated with indicated concentrations of bemcentinib for 1 hour, infected with MHV (MOI = 0.001) for 24 hours and viral load assessed by RT-qPCR. **C)** As in B, MHV infection of peritoneal macrophages (MOI =0.001) treated with indicated concentrations of bemcentinib. Data shown are representative of 3 independent experiments. Data represented as means ± SEM. Student's t-test; asterisks represent $p < 0.05$.

**Supplemental Figure 8:** Author contributions, as defined by CRediT (Contributor Roles Taxonomy).

| | Conceptualization | Methodology | Software | Validation | Formal Analysis | Investigation | Resources | Data Curation | Writing- Original Draft | Writing - Review and editing | Visualization | Supervision | Project Administration | Funding Acquisition |
|---|---|---|---|---|---|---|---|---|---|---|---|---|---|---|
| **D.B.** | ■ | ■ | | ■ | ■ | ■ | | ■ | ■ | ■ | ■ | | ■ | |
| **H.V.E.** | ■ | ■ | | ■ | ■ | ■ | | ■ | | ■ | ■ | | | |
| N.R. | | | | | ■ | ■ | | | | | | | | |
| K.J.R. | ■ | | | | ■ | ■ | | | | ■ | | | | |
| M.B. | | | | | | ■ | | | | | | | | |
| J.A.A.B | | ■ | | | | ■ | | | | ■ | | | | |
| R.A.R.C. | | | | | | ■ | | | | | | | | |
| B.G. | | | | | | | ■ | | | | | | | |
| T.S. | | | ■ | | ■ | | | ■ | | | | | | |
| E.C. | | | | | | ■ | | | | | | | | |
| D.M. | | ■ | | | ■ | ■ | ■ | | | | | | | |
| G.G. | | | | | | | | | | ■ | | | | |
| H.H. | | ■ | | | ■ | ■ | ■ | | | | | | | |
| J.M. | ■ | | | | | | ■ | | | | | | | |
| J.B.L. | ■ | ■ | | | | ■ | ■ | | | | | ■ | | |
| **W.J.M.** | ■ | ■ | | ■ | ■ | ■ | ■ | ■ | ■ | ■ | | ■ | ■ | ■ |
